# Tregs constrain CD8^+^ T cell priming required for curative intratumorally anchored anti-4-1BB immunotherapy

**DOI:** 10.1101/2023.01.30.526116

**Authors:** Joseph R. Palmeri, Brianna M. Lax, Joshua M. Peters, Lauren Duhamel, Jordan A. Stinson, Luciano Santollani, Emi A. Lutz, William Pinney, Bryan D. Bryson, K. Dane Wittrup

## Abstract

Although co-stimulation of T cells with agonist antibodies targeting 4-1BB (CD137) improves antitumor immune responses in preclinical studies, clinical development has been hampered by on-target, off-tumor toxicity. Here, we report the development of a tumor-anchored α4-1BB agonist (α4-1BB-LAIR), which consists of an α4-1BB antibody fused to the collagen binding protein LAIR. While combination treatment with an antitumor antibody (TA99) displayed only modest efficacy, simultaneous depletion of CD4^+^ T cells boosted cure rates to over 90% of mice. We elucidated two mechanisms of action for this synergy: αCD4 eliminated tumor draining lymph node Tregs, enhancing priming and activation of CD8^+^ T cells, and TA99 + α4-1BB-LAIR supported the cytotoxic program of these newly primed CD8^+^ T cells within the tumor microenvironment. Replacement of αCD4 with αCTLA-4, a clinically approved antibody that enhances T cell priming, produced equivalent cure rates while additionally generating robust immunological memory against secondary tumor rechallenge.

**One Sentence Summary:** Inhibition of nodal Tregs enhances CD8^+^ T cell priming, improving antitumor responses to collagen-anchored α4-1BB combination therapy.

## INTRODUCTION

The use of monoclonal antibodies to perturb immune cell signaling networks and improve anti-cancer immune responses has gained increased attention in recent years (*1*). Checkpoint blockade therapy with antagonistic antibodies is safe and efficacious, but agonistic antibodies against targets such as 4-1BB, OX40, GITR, and ICOS have proven to exhibit impractically narrow therapeutic windows due to on-target, off-tumor toxicity (*2–5*).

4-1BB (also known as CD137 or TNFRSF9) is expressed primarily on activated CD8^+^ and CD4^+^ T cells, including CD4^+^ regulatory T cells (Tregs), and natural killer (NK) cells, and is a promising target for agonist antibodies (*6–10*). Signaling through 4-1BB in CD8^+^ T cells leads to proliferation, enhanced survival, cytokine production, improved memory formation, and altered metabolism (*11–15*). Treating mice with agonist α4-1BB antibodies as a monotherapy or in combination therapies is highly efficacious in several preclinical mouse cancer models (*16, 17*). However, toxicity has hampered clinical translation of such antibodies, with lethal liver toxicities reported in early phase 2 trials of Urelumab, the first α4-1BB agonistic antibody to enter the clinic (*18*). At reduced doses which do not elicit dose limiting toxicities (DLTs), little to no clinical efficacy has been reported (*18*). Utomilumab, the second α4-1BB agonist to enter the clinic, is well tolerated but is a much weaker agonist and has little clinical activity (*19, 20*). Given the difficulty of uncoupling toxicity from clinical activity with systemically administered agonists, recent development around this target has focused on engineering antibodies with tumor specific activity (*21*). This includes several bispecific antibodies, with one arm targeting 4-1BB and the other targeting either tumor specific antigens or PD-L1, α4-1BB antibodies that bind only in tumor specific niches, such as high ATP concentrations, or pro-drug α4-1BB antibodies where the binding domain of the antibody is shielded by a peptide “mask” that is cleaved by tumor specific proteases (*22–26*).

Alternatively, our group and others have demonstrated the utility of using collagen binding strategies to anchor immunotherapy payloads to the tumor microenvironment (*27–35*). Collagen is a desirable target for localization due to its abundance in the tumor microenvironment (TME) (*36*). By directly fusing collagen binding domains to cytokines and chemokines or chemical conjugation of collagen binding peptides to αCTLA-4 and αCD40 antibodies, intratumoral administration of these therapeutic payloads results in prolonged tumor retention, enhanced efficacy, and reduced systemic toxicity.

In this work, we developed a locally retained collagen-anchored α4-1BB agonist, termed α4-1BB-LAIR, by fusing an α4-1BB agonist to the ectodomain of an endogenous collagen binding protein, Leukocyte Associated Immunoglobulin Like Receptor 1 (LAIR1). Tested in combination with an antitumor antibody, TA99, in a fully syngeneic and poorly immunogenic B16F10 murine melanoma model, this combination exhibited little efficacy. Intriguingly, depletion of CD4^+^ T cells led to long term durable cures in >90% of TA99- + α4-1BB-LAIR-treated animals. However, nearly all of these mice were unable to control a secondary tumor rechallenge. We hypothesized that depletion of Tregs, which comprise a subset of CD4^+^ T cells, was driving this synergy. Tregs are immunosuppressive CD4^+^ T cells that express the transcription factor forkhead box P3 (Foxp3) and are critical to maintaining homeostasis and preventing autoimmunity (*37, 38*). Indeed, *Foxp3^−/−^* mice die at a young age from severe lymphoproliferative disease, systemic depletion of Tregs in adult mice leads to rapid lethal autoimmunity, and *Foxp3* mutations in humans cause severe immune dysregulation (*39–42*). Although Tregs play a critical role in curbing autoreactive T cells, they also constrict productive antitumor immune responses through a variety of mechanisms and at various stages of the tumor-immunity cycle (*43, 44*).

Using flow cytometry and bulk-RNA sequencing, we probed the immunological mechanism of this synergy and found that CD4^+^ T cell depletion led to an enhanced activation state in the tumor draining lymph node (TdLN), leading to an influx of newly primed CD8^+^ T cells into the tumor. Local remodeling of the tumor microenvironment by TA99 and α4-1BB-LAIR enhanced the cytotoxicity of these newly primed T cells, leading to tumor cell death and eventual complete tumor regression. Using a Foxp3-DTR mouse model, which allows for selective depletion of Tregs, we confirmed that Treg depletion alone was sufficient for this synergy. Finally, we demonstrated that CD4^+^ T cell depletion can be replaced with a more clinically relevant agent known to enhance CD8^+^ T cell priming, αCTLA-4, without compromising efficacy. This combination of TA99 + α4-1BB-LAIR + αCTLA-4 also resulted in formation of robust immunological memory, enabling rejection of a secondary tumor rechallenge. This work suggests that locally retained 4-1BB agonist and antitumor antibody therapy can be highly efficacious when combined with modalities that enhance T cell priming, which can be restrained by TdLN Tregs.

## RESULTS

### TA99 + α4-1BB-LAIR synergizes robustly with CD4 compartment depletion

In order to develop a tumor-localized 4-1BB agonist, we leveraged a collagen anchoring strategy previously validated by our lab and others. We recombinantly expressed an α4-1BB antibody (clone LOB12.3, Table S1) as a C-terminal fusion with the ectodomain of murine LAIR1, an endogenous immune cell inhibitory receptor that naturally binds collagen (*45–47*). We verified that α4-1BB-LAIR expresses without aggregation (Fig. S1A), is able to bind plate bound collagen I via ELISA (Fig. S1B), and that binding to cell-surface expressed 4-1BB is unaffected.

To validate intratumoral retention without confounding target-mediated drug disposition (TMDD) effects, we also generated an αFITC-LAIR control antibody as this antibody has no known murine target (Table S1). Fluorescently labeled αFITC-LAIR administered intratumorally was preferentially retained in the tumor over unanchored αFITC antibody when measured longitudinally with IVIS (Fig. S1D-E).

α4-1BB agonist Urelumab is being clinically tested in combination with antitumor antibodies Rituximab, Cetuximab, and Elotuzumab which target CD20, EGFR, and SLAMF7, respectively (NCT01775631, NCT02110082, NCT02252263). Preliminary data has not been encouraging, with early reports from the Rituximab combination suggesting that Rituximab + Urelumab is no more efficacious than Rituximab monotherapy (*48*). We sought to evaluate if collagen-anchoring would improve the efficacy of combination α4-1BB agonist + antitumor antibody therapy. Mice were inoculated with B16F10 melanoma flank tumors and treated systemically (intraperitoneally, or i.p.) with TA99, an antitumor antibody that binds to Trp1 expressed on the surface of B16F10 cells, followed by intratumorally (i.t.) administered α4-1BB-LAIR one day later for a total of 4 weekly cycles (This combination of TA99 + α4-1BB-LAIR is referred to collectively as the treatment, or “Tx”, henceforth, Fig. 1A). Although this combination led to a statistically significant growth delay compared to PBS treated mice, nearly all mice eventually succumbed to their tumor burden, with only ~5% of mice achieving a complete response (CR, defined as no palpable tumor at day 100) (Fig. 1B). Notably, this growth delay was only slightly better than the individual components of Tx, although it was the only therapy with any complete responders (Fig. S2A).

**Figure 1.**
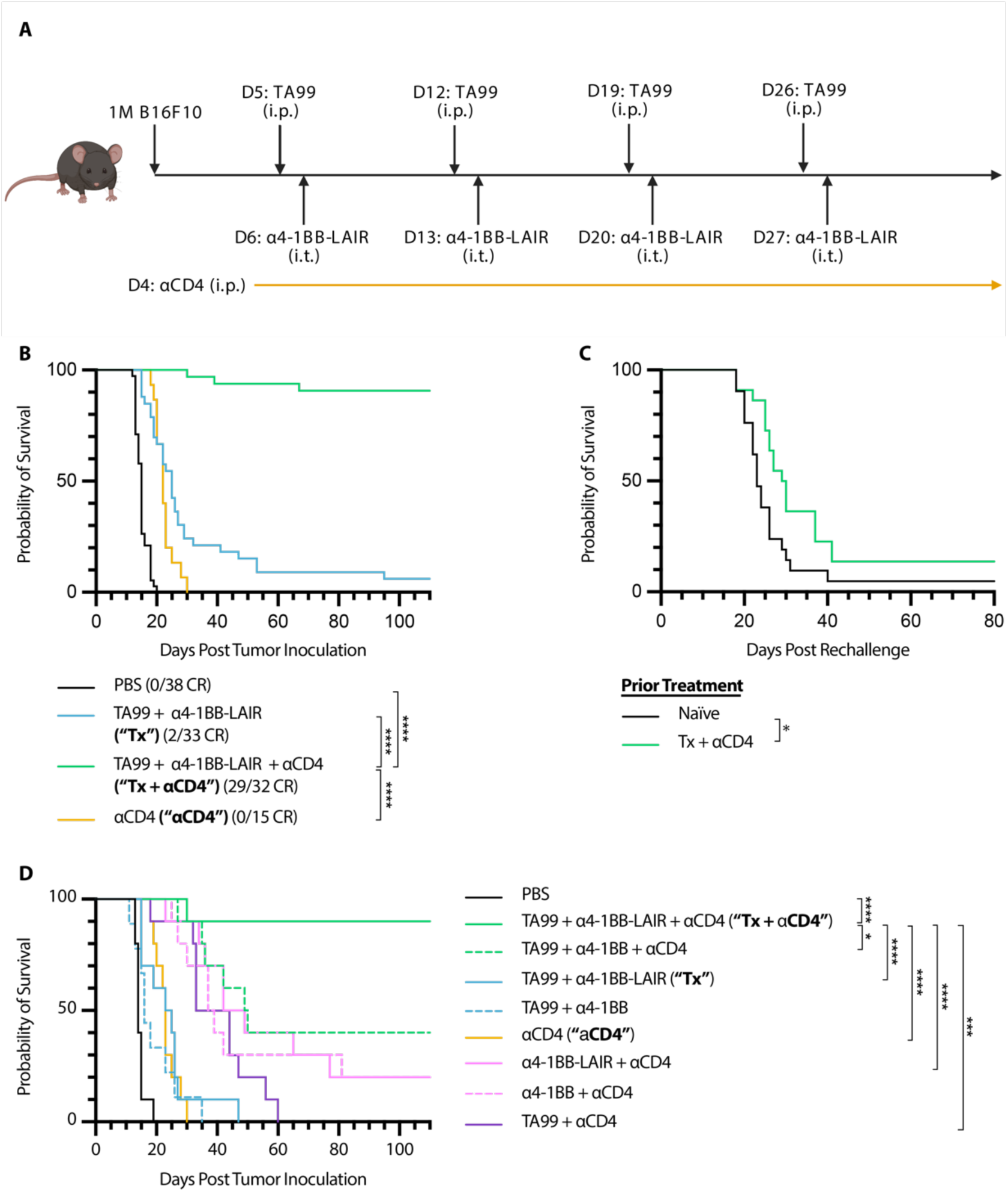
TA99 + α4-1BB-LAIR synergizes robustly with CD4^+^ T cell depletion. Mice were inoculated with 1 x 10^6^ B16F10 cells on day 0. **(A)** Treatment schedule of TA99 + α4-1BB-LAIR + αCD4. Mice were treated with 200 µg of TA99 (i.p.) on days 5, 12, 19, and 26, treated with 36.1 µg α4-1BB-LAIR (i.t.) on days 6, 13, 20, and 27 (molar equivalent to 30 µg α4-1BB), and treated with 400 µg αCD4 (i.p.) every 3 days starting 1 day before the first dose of TA99 and ending one week after the last dose of α4-1BB-LAIR (days 4 to 34). **(B)** Aggregate survival of mice treated with PBS (n = 38), TA99 + α4-1BB-LAIR (“**Tx**”) (n = 33), TA99 + α4-1BB-LAIR + αCD4 (“**Tx + αCD4**”) (n = 32), or αCD4 (n = 15) (eight independent studies). **(C)** Survival of complete responders to Tx + αCD4 re-challenged on the contralateral flank >100 days after primary tumor inoculation. **(D)** Overall survival of mice treated with indicated combination variants, demonstrating all components are necessary for maximum efficacy (n = 9-10, two independent experiments). Survival was compared using log-rank Mantel-Cox test. \**P* < 0.05, \*\**P* < 0.01, \*\*\**P* < 0.001, \*\*\*\**P* < 0.0001.

In an effort to improve this combination therapy, we explored which cell types were critical for response. Surprisingly, we observed that when we also treated these mice with an αCD4 antibody that depletes the entire CD4^+^ T cell compartment, the complete response rate of TA99 + α4-1BB-LAIR improved dramatically, with >90% of mice achieving a complete response (Fig. 1B). However, when long-term survivors were rechallenged on the contralateral flank >100 days after initial tumor inoculation, nearly all mice succumbed to these secondary tumors (Fig. 1C). This was indicative of the inability of these mice to develop robust immune memory to B16F10 tumor cells, likely resulting from the depletion of CD4^+^ effector T cells.

Growth delay with systemically administered αCD4 and α4-1BB has been reported previously, but we find that our specific components were necessary to achieve maximum efficacy, including TA99 (*P* = .0032) and, notably, retention via collagen anchoring (*P* = .0289) (Fig. 1D) (*49*). Consistent with other preclinical reports with this α4-1BB antibody clone, no signs of toxicity were observed for the full therapeutic combination with or without collagen anchoring (Fig. S2B) (*50*). Although αCD4 drastically improved the efficacy of Tx, the lack of immune memory formation and low translational potential of long term αCD4 treatment motivated us to understand the mechanism of this synergy and ultimately develop alternative clinically relevant synergistic combinations.

### αCD4 improves priming in the TdLN

We investigated the chemokine/cytokine profile of the TME following treatments with PBS, Tx, Tx + αCD4, or αCD4 both 3 and 6 days after α4-1BB-LAIR administration. We dissociated tumors and analyzed the cytokine and chemokine milieu using a multiplexed flow cytometry-based ELISA assay. Although we observed general increases in inflammatory cytokines and chemokines in all treatment groups, only GM-CSF was specifically upregulated in the Tx + αCD4 group when compared to Tx or αCD4 alone (Fig. S3A). However, neutralization of this cytokine did not abrogate therapeutic efficacy of Tx + αCD4, indicating that this spike in GM-CSF was dispensable for therapeutic efficacy (Fig. S3B).

We then used flow cytometry to analyze the tumors and tumor draining lymph nodes (TdLNs) of mice treated with Tx, Tx + αCD4, or αCD4 again 3 and 6 days after the first α4-1BB-LAIR treatment. As expected, we observed complete depletion of total CD4^+^ T cells and Tregs (defined as Foxp3^+^ CD25^+^ CD4^+^ T cells) in the tumor (Fig. 2A) and TdLN (Fig. 2B) in both the Tx + αCD4 and the αCD4 groups.

**Figure 2.**
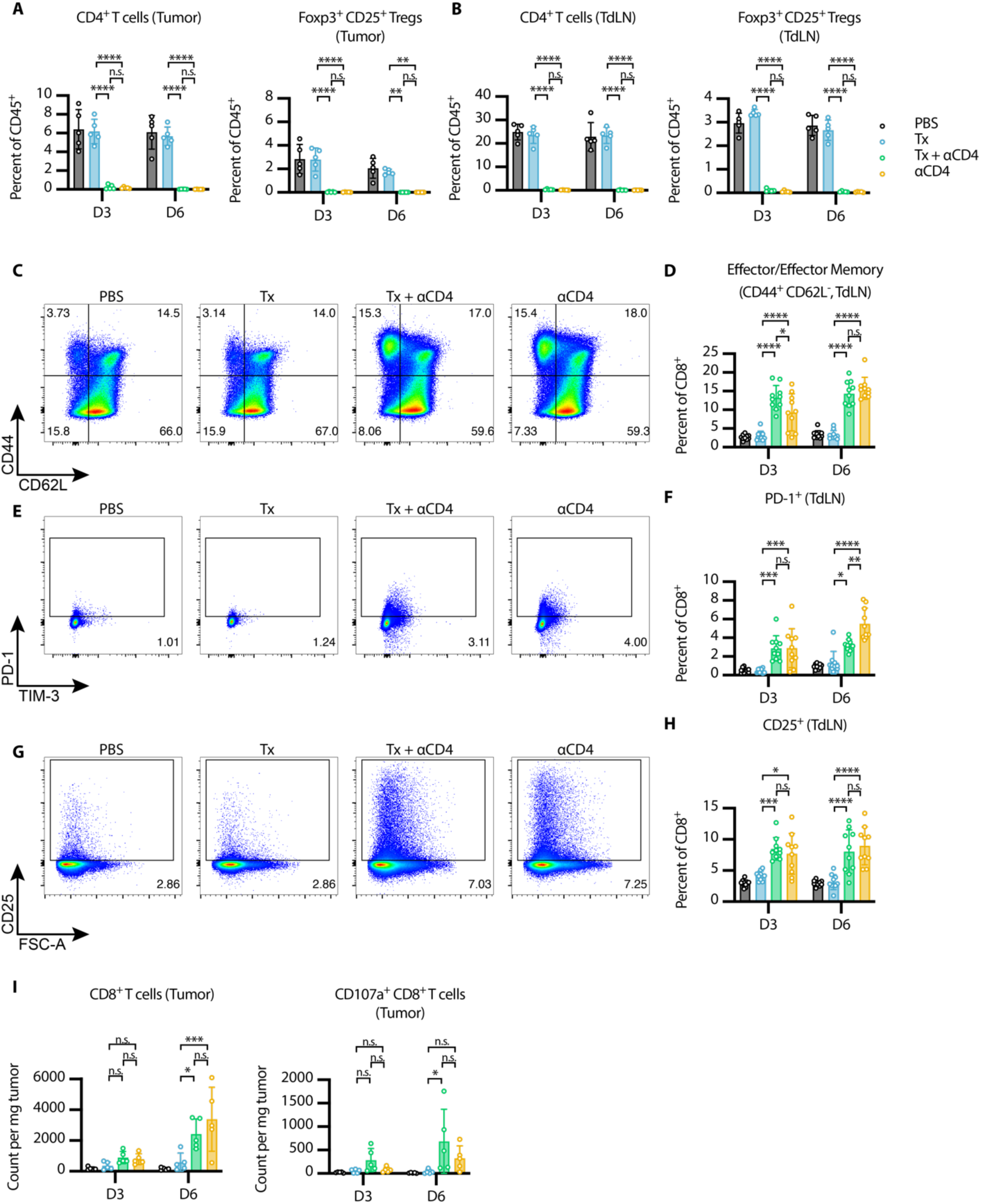
αCD4 leads to new wave of CD8^+^ T cell priming and Tx supports cytotoxicity of these cells in the tumor. **(A-B)** Flow cytometry quantification (mean±SD) of CD4^+^ T cells (gated on single cell/live/CD45^+^/CD3^+^NK1.1^−^/CD4^+^) and Tregs (gated on single cell/live/CD45^+^/CD3^+^NK1.1^−^/CD4^+^/Foxp3^+^CD25^+^) in the **(A)** tumor and **(B)** TdLN 3 and 6 days after first α4-1BB-LAIR treatment (n = 5). **(C)** Representative gating on CD44 and CD62L to define effector/effector memory CD8^+^ T cells in TdLN 6 days after first α4-1BB-LAIR treatment and **(D)** quantification (mean±SD) of these cell populations 3 and 6 days after first α4-1BB-LAIR treatment (gated on single cell/live/CD45^+^/CD3^+^NK1.1^−^/CD8^+^/CD44^+^CD62L^−^, n = 10, two independent experiments). **(E)** Representative gating of PD-1^+^ CD8^+^ T cells 6 days after first α4-1BB-LAIR treatment and **(F)** quantification (mean±SD) of these cell populations 3 and 6 days after first α4-1BB-LAIR treatment (gated on single cell/live/CD45^+^/CD3^+^NK1.1^−^/CD8^+^/PD-1^+^, n = 10, two independent experiments) **(G)** Representative gating of CD25^+^ CD8^+^ T cells 6 days after first α4-1BB-LAIR treatment and **(H)** quantification (mean±SD) of these cell populations 3 and 6 days after first α4-1BB-LAIR treatment (gated on single cell/live/CD45^+^/CD3^+^NK1.1^−^/CD8^+^/CD25^+^, n = 10, two independent experiments). **(I)** Flow cytometry quantification (mean±SD) of CD8^+^ T cells (left) and CD107a^+^ CD8^+^ T cells (right) (gated on single cell/live/CD45^+^/CD3^+^NK1.1^−^/CD8^+^) in the tumor 3 and 6 days after first α4-1BB-LAIR treatment (n = 5). Flow cytometry data was compared using two-way ANOVA with Tukey’s multiple hypothesis testing correction. \**P* < 0.05, \*\**P* < 0.01, \*\*\**P* < 0.001, \*\*\*\**P* < 0.0001.

Using CD44 and CD62L gating, we divided CD8^+^ T cells in the TdLN into naive (CD44^−^ CD62L^+^), effector/effector memory (CD44^+^ CD62L^−^), and central memory (CD44^+^ CD62L^+^) phenotypes. At both time points, we observed a shift of the CD8^+^ T cell population towards an effector/effector memory phenotype in the Tx + αCD4 and αCD4 groups (Fig. 2C-D). Additionally, we observed increases in both PD-1^+^ CD8^+^ T cells (Fig. 2E-F) and CD25^+^ CD8^+^ T cells (Fig. 2G-H), at both time points in the Tx + αCD4 and αCD4 groups, both of which are markers of recently activated CD8^+^ T cells in lymphoid tissue. The magnitude of these changes was equivalent between the Tx + αCD4 and αCD4 groups, indicating that the αCD4 antibody component was driving these changes to the TdLN.

6 days following treatment with either Tx + αCD4 or αCD4, we observed increased CD8^+^ T cells infiltrating the tumor (Fig. 2I), which is in agreement with the enhanced activation state observed in the TdLN (Fig. 2E-H). This result is consistent with previous preclinical and clinical studies that have shown treatment with αCD4 can enhance T cell priming, leading to increased numbers of tumor reactive CD8^+^ T cells (*51–53*). However, only in the Tx + αCD4 group, when compared to PBS or Tx alone, do we observe an increase in degranulating CD107a^+^ CD8^+^ cytotoxic T cells (Fig. 2I, Fig. S4A). No major differences in 4-1BB expression on CD8+ T cells were detected in the tumor and only minor increases were seen on CD8^+^ T cells in the TdLN in Tx + αCD4 and αCD4 treated mice (Fig. S4B-D). These data suggest that αCD4 therapy, independent of Tx, induces *de novo* priming in the TdLN, leading to more CD8^+^ T cell infiltration in the tumor. However, we hypothesized that Tx supports these newly primed cells and maintains their cytotoxic phenotype within the tumor, leading to eventual tumor regression.

### TdLN has increased proliferation and T cell gene signatures by Bulk-RNA sequencing

To further interrogate immunological changes to the TdLN and tumor in an unbiased holistic manner, we performed bulk RNA-sequencing on CD45^+^ cells from TdLN samples from mice treated with PBS, Tx, Tx + αCD4, or αCD4 3 and 6 days following α4-1BB-LAIR administration. We generated a UMAP plot of the TdLN samples and found that, at the bulk transcript level, large differences between samples were apparent only at the later time point (Fig. 3A). Additionally, sample clustering at this later time point was driven entirely by αCD4, with the αCD4 and Tx + αCD4 samples clustering separately from the PBS and Tx samples. In fact, we observed almost no differentially expressed genes (DEGs) in the TdLN when comparing Tx + αCD4 versus αCD4 or Tx versus PBS treated samples (Fig. 3B), indicating that Tx alone had no appreciable change on the transcriptional program in the TdLN.

**Figure 3.**
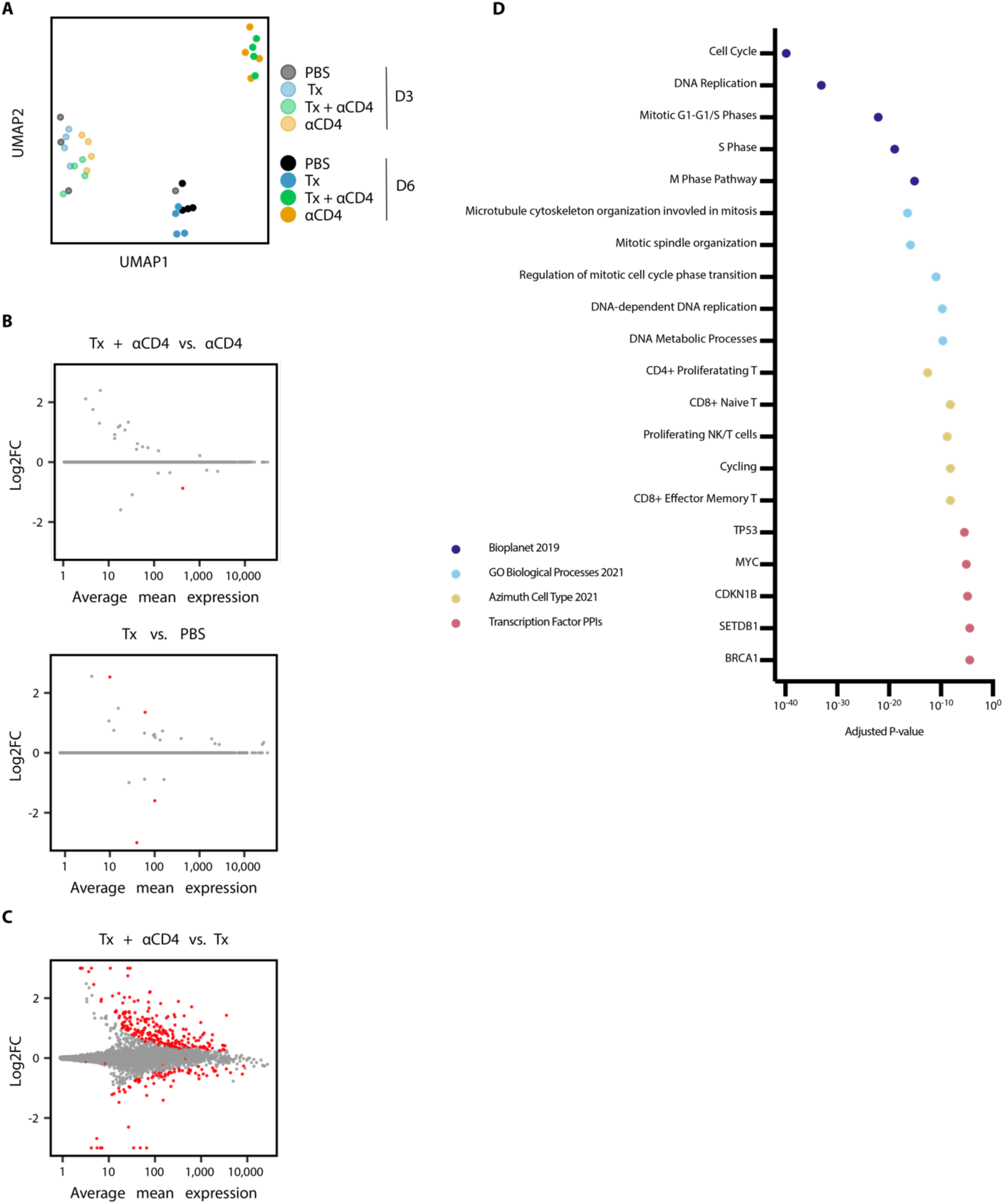
αCD4 drives proliferation in the TdLN. **(A)** UMAP plot of TdLN transcriptomes (n = 4 per group) **(B)** Differential expression testing of Tx + αCD4 vs. αCD4 and Tx vs. PBS TdLN samples 6 days after first α4-1BB-LAIR treatment, with statistically significant hits highlighted in red (FDR ≤ 5%). **(C)** Differential expression testing of Tx + αCD4 vs. Tx TdLN samples 6 days after to first α4-1BB-LAIR treatment, with statistically significant hits highlighted in red (FDR ≤ 5%). **(D)** Pathway enrichment analysis of upregulated DEGs identified in **(C)**.

To assess what changes αCD4 drove in the TdLN, we examined DEGs between Tx + αCD4 and Tx treated samples (Fig. 3B). We found 247 upregulated genes and 82 downregulated genes (FDR ≦ 5%, Fig. 3C). We used enrichR to determine which pathways these upregulated DEGs were enriched in (*54–56*). Upregulated genes belonged to pathways involving cell cycling, DNA replication, and Myc related genes, indicative of a highly proliferative state in the TdLN. They were also enriched for both cycling and CD8^+^ T cell states (Fig. 3D). Overall, the TdLN transcriptional data demonstrated that 1) changes to the TdLN resulted from αCD4 treatment, independent of Tx, and 2) these changes led to enhanced proliferation and T cell activation in the TdLN.

### Tx + αCD4 leads to cytotoxic CD8^+^ T cell program in the tumor

We similarly used bulk-RNA sequencing to examine immune cell gene expression programs within the tumor. We performed hierarchical clustering of the tumor samples while also independently clustering all significant DEGs (with a log 2-fold change ≥2 or ≤-2 and *p*-adj ≤ 0.05) using *k*-means clustering. This clustering identified 10 distinct gene clusters of co-expressed genes. Samples clustered imperfectly by treatment type, with two of the three Tx + αCD4 day 6 samples showing distinct transcriptional programs (Fig. 4A-B, Fig. S5). These two samples had the smallest tumor size at time of necropsy, indicating they were already robustly responding to therapy at this time point. We next performed pathway enrichment analysis on the individual gene clusters. Of particular interest were cluster 1 and 2 (and to a lesser extent cluster 4), which were upregulated specifically in the Tx + αCD4 groups, and cluster 3, which contains genes upregulated in both the Tx + αCD4 and Tx groups and representing a Tx-specific transcriptional program (Fig. 4B). These clusters are enriched for a range of GO terms associated with productive cellular immune responses (regulation of T cell activation, alpha-beta T cell activation, lymphocyte mediated immunity, etc.). However, only clusters 1 and 2 were enriched for genes associated with interferon gamma production, suggesting that Tx alone is not sufficient to drive IFNγ production (Fig. 4C). Notably, because Tx + αCD4 and αCD4 drive similar levels of increased CD8+ T cell counts (Fig. 2F), but cytotoxic genes are only enriched in Tx + αCD4, we can conclude that this cytotoxic signature is not an artifact of increased CD8+ T cell counts. Cluster 7, which is highly expressed in PBS samples, contained genes enriched for, among others, pigmentation gene programs, likely representing increased CD45^−^ tumor cells in this sample (Fig. 4C).

**Figure 4.**
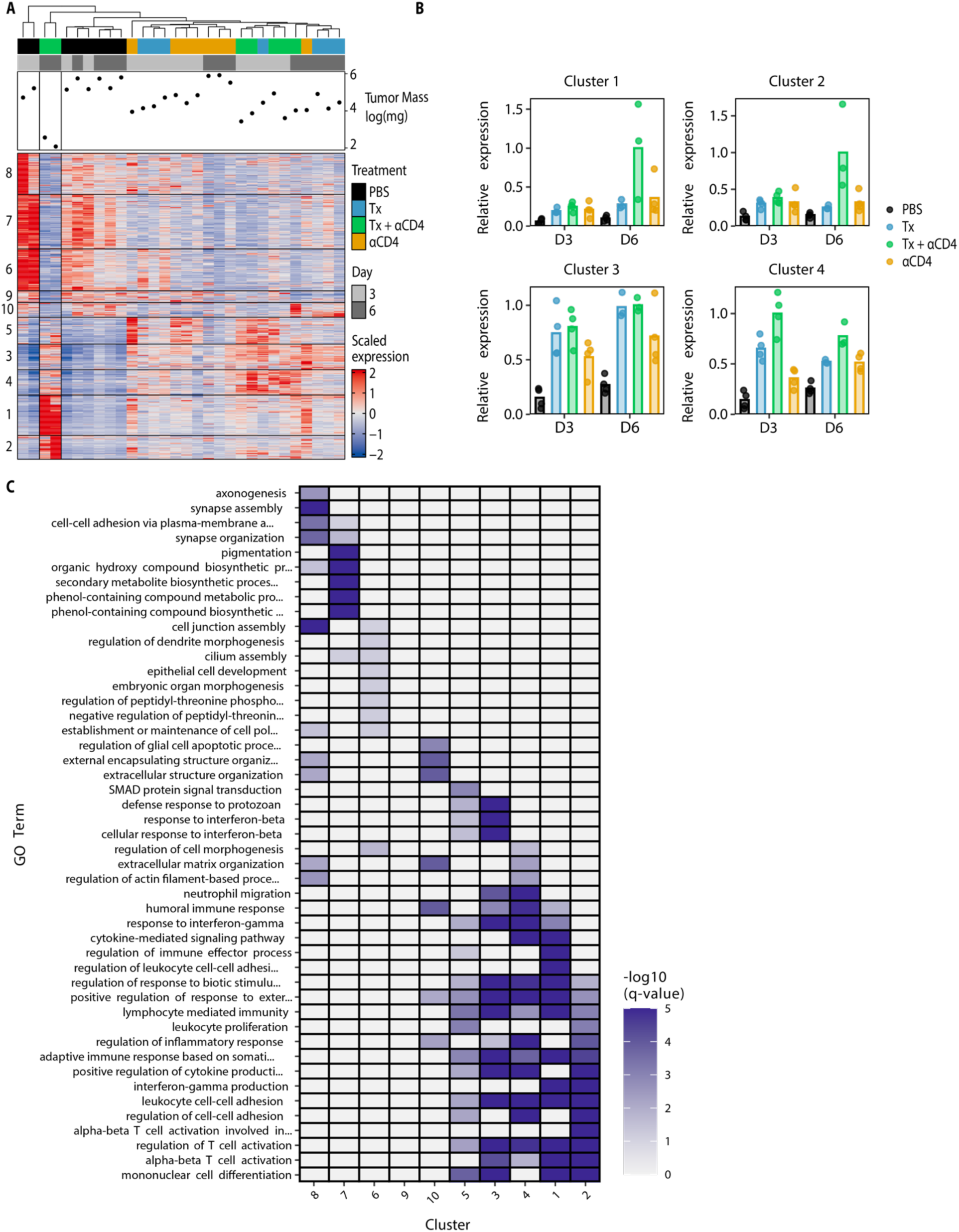
Tx + αCD4 upregulated gene clusters enriched for CD8^+^ effector programs. **(A)** Heatmap of *k*-means clustered DEGs (absolute value lfc ≥ 2, FDR ≤ 10%) and tumor samples hierarchically clustered (n = 3-4) **(B)** Normalized expression of select individual gene clusters identified in **(A)** for each experimental condition. **(C)** Pathway enrichment analysis for each gene cluster identified in **(A).**

To further assess changes to the tumor microenvironment, we looked at DEGs between Tx + αCD4 tumor samples 3 and 6 days following α4-1BB-LAIR. 63 genes were upregulated, and 43 genes downregulated between these two time points (FDR ≤ 5%, Fig. 5A). We used the upregulated DEGs to establish a “response” signature for Tx + αCD4. We then asked if this gene signature was expressed in any other treatment conditions/time points. Indeed, this signature was highly expressed only in the Tx + αCD4 late time point, indicating this was a bona fide response signature unique to Tx + αCD4 treated mice (Fig. 5B). We then performed pathway enrichment analysis to determine what pathways these genes were associated with. Confirming our previous flow data, we saw effector and effector memory T cell signatures. Additionally, we saw genes associated with TCR signaling, interleukin-2 (IL-2) signaling and Stat5a activity (Fig. 5C). Recent literature has highlighted a role for IL-2, or more broadly Stat5a activity, in amplifying T cell populations that drive responses to checkpoint blockade (*57–59*). Taken together, the tumor transcriptional data support the notion that Tx + αCD4 drives a robust cytotoxic T cell program leading to tumor rejection.

**Figure 5.**
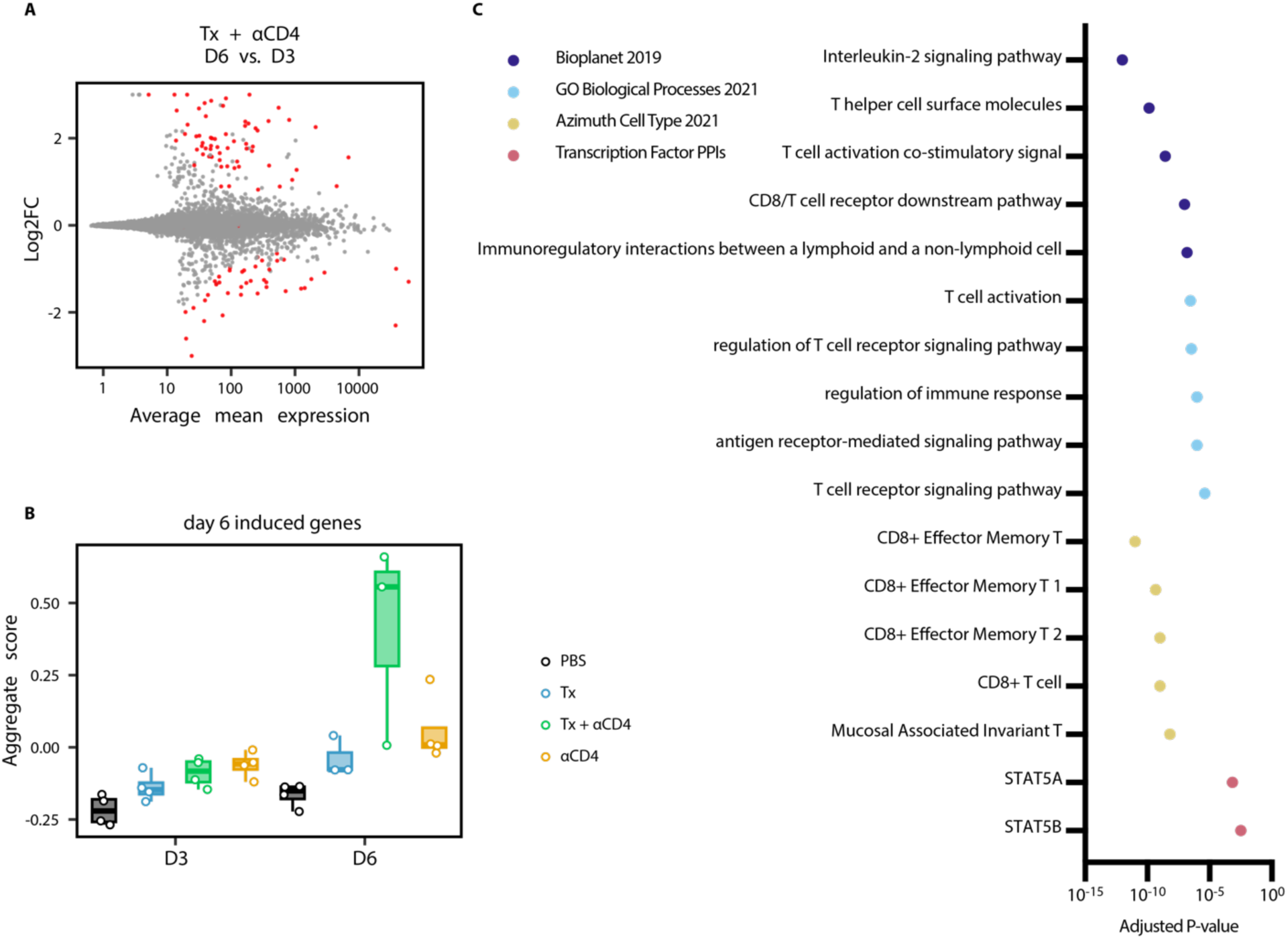
Tx + αCD4 associated with cytotoxic T cell signature in the tumor. **(A)** Differential expression testing of Tx + αCD4 on day 3 vs. day 6 tumor samples relative to first α4-1BB-LAIR treatment, with statistically significant hits highlighted in red (FDR ≤ 5%). **(B)** Average expression level of significantly upregulated DEGs identified in **(A)** across all treatment groups. **(C)** Pathway enrichment analysis of upregulated DEGs identified in **(A)**.

### Treg depletion results in equivalent efficacy as whole CD4 compartment depletion

We hypothesized that Treg depletion was the primary functional consequence of αCD4 therapy, and that Treg specific elimination would lead to similar efficacy in combination with Tx. To test this hypothesis, we turned to Foxp3-DTR mice, which express the diphtheria toxin receptor (DTR) and GFP under control of the Foxp3 promoter. In these mice, all Foxp3^+^ cells are also DTR^+^, and thus susceptible to diphtheria toxin (DT) mediated cell death. Systemic administration of DT to these mice leads to rapid and complete depletion of nearly all Foxp3^+^ Tregs. However, with repeat dosing these mice succumb to lethal autoimmunity within 10-20 days of DT administration (*39*). In order to facilitate long term depletion of Tregs in the tumor and TdLN without inducing lethal autoimmunity, we developed a low dose, intratumoral diphtheria regimen. Every other day intratumoral dosing of 75ng or 125ng of DT depleted tumor and TdLN to similar levels as 1µg of systemically administered DT, with reduced impacts on splenic Treg populations (Fig. 6A, S6A). Additionally, we did not observe signs of toxicity, as measured by weight loss, with intratumoral low dose DT, while mice receiving systemic DT showed trends of weight loss at time of euthanasia (Fig. 6B, S6B). Thus, we felt confident that low dose intratumoral DT was a safe and effective model system to achieve long term intratumoral and intranodal Treg depletion

**Figure 6.**
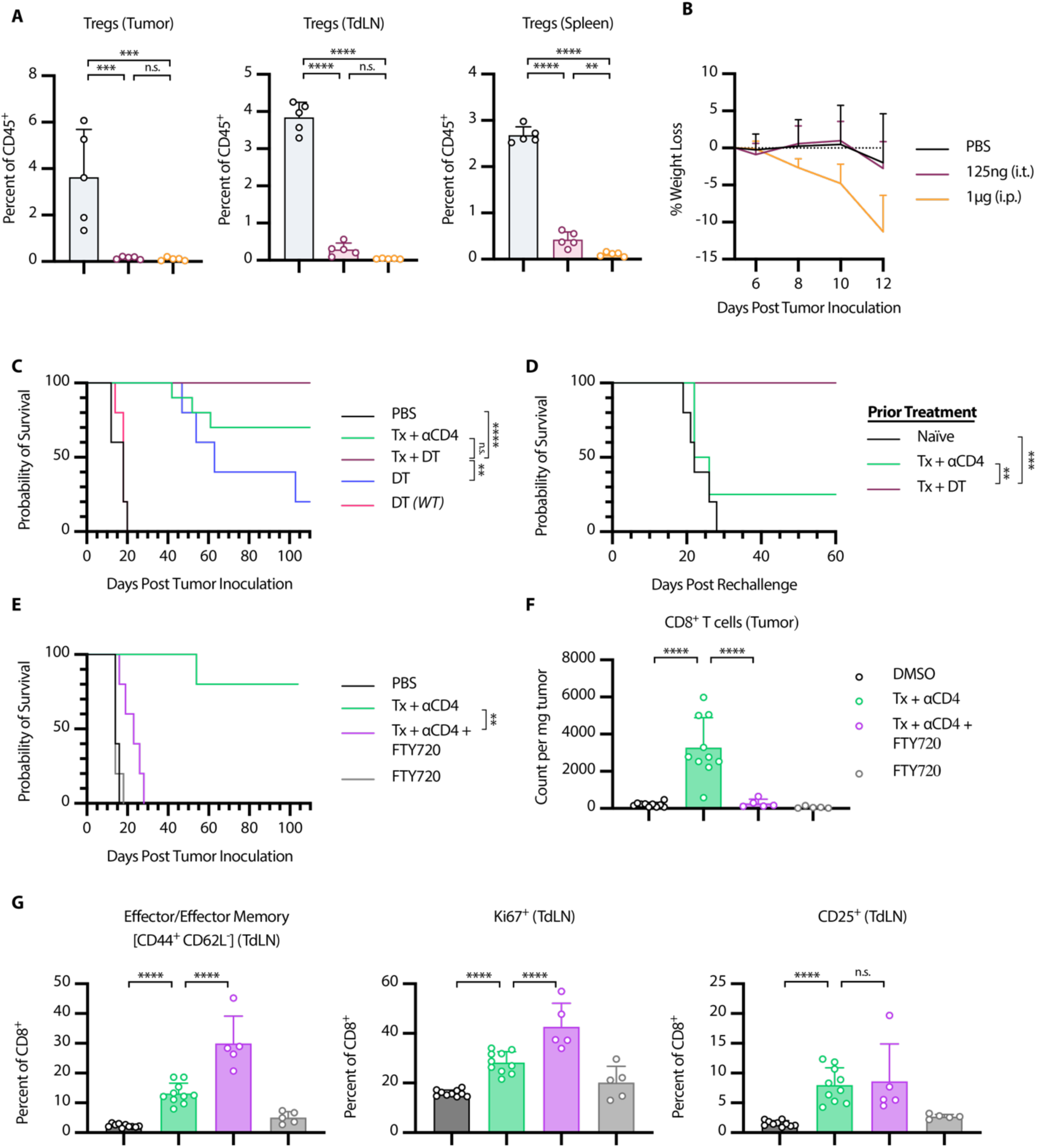
Tx + αCD4 is Treg dependent and requires *de novo* priming for efficacy. **(A-D)** Foxp3-DTR Mice were inoculated with 1 x 10^6^ B16F10 cells on day 0. **(A)** Mice were treated on days 6, 8, and 10 with either 125 ng DT (i.t.) or 1 µg DT (i.p.). Flow cytometry quantification (mean±SD) of Tregs in tumor, TdLN, or spleen on day 12 (gated on single cell/live/CD45^+^/CD3^+^NK1.1^−^/CD4^+^/GFP(*Foxp3*)^+^, n = 5). **(B)** Weight loss (mean+SD) of mice from **(A)**. **(C)** Survival of Foxp3-DTR mice treated with PBS (n = 5), Tx + αCD4 (n = 10), Tx + DT (n = 10), DT (n = 5), WT mice treated with DT (n = 5). Mice were treated with the same relative dose/dose schedule as in Fig. 1A, but treatment initiation was delayed two days. DT treated mice received 125 ng DT (i.t.) every other day from day 6 to day 36. **(D)** Survival of complete responders to Tx + αCD4 or Tx + DT re-challenged on the contralateral flank >100 days after primary tumor inoculation. **(E-G)** WT mice were inoculated with 1 x 10^6^ B16F10 cells on day 0. **(E)** Overall survival of mice treated with PBS/DMSO (n = 5), Tx + αCD4 (n = 5), Tx + αCD4 + FTY720 (n = 5), or FTY720 (n = 5). Mice were treated with the same relative dose/dose schedule as in Fig. 1A, but treatment initiation was delayed two days. Mice were treated with 30 µg of FTY720 (i.p.) every other day from days 6 to 36. **(F)** Flow cytometry quantification (mean±SD) of CD8^+^ T cell counts in tumor 6 days after first α4-1BB-LAIR treatment. (gated on single cell/Live/CD45^+^/CD3^+^NK1.1^−^/CD8^+^, n = 5-10, two independent experiments). **(G)** Flow cytometry quantification (mean±SD) of effector/effector memory (CD44^+^ CD62L^−^), CD25^+^, and Ki67^+^ CD8^+^ T cells in the TdLN 6 days after α4-1BB-LAIR treatment (gated on single cell/live/CD45^+^/CD3^+^NK1.1^−^/CD8^+^, n = 5-10, two independent experiments). Flow cytometry data was compared using one-way ANOVA with Tukey’s multiple hypothesis testing correction. Survival was compared using log-rank Mantel-Cox test. \**P* < 0.05, \*\**P* < 0.01, \*\*\**P* < 0.001, \*\*\*\**P* < 0.0001.

B16F10 tumor-bearing Foxp3-DTR mice were treated with Tx + αCD4, Tx + DT, or DT alone. To allow for lesions of sufficient size for intratumoral administration of DT, the absolute timing of therapy administration was delayed two days for all groups (such that DT and αCD4 were initiated on day 6). Mice receiving Tx + DT responded equally as well as mice receiving Tx + αCD4, with a trend (but not statistically significant) towards a higher complete response rate in the Tx + DT group (Fig. 6C). Interestingly, DT on its own also resulted in significant growth delay, but ultimately almost all mice succumbed to their tumor burden. To confirm that the effect of DT was purely a result of Treg depletion, we treated WT mice with DT, which resulted in no different growth kinetics over PBS treated mice. No signs of toxicity, as assessed by weight loss, were observed throughout the course of treatment (Fig. S6C). A previously published study demonstrated that transient DT given with systemic α4-1BB agonist therapy led to severe immune related adverse events (irAEs) in MC38 tumor bearing mice, further highlighting the advantages of our collagen-anchored α4-1BB agonists (*60*). Notably, when cured mice were rechallenged >100 days after their primary tumor inoculation, the majority of the Tx + αCD4 cured mice did not reject rechallenge, consistent with previous results, while 100% of mice cured with Tx + DT rejected this rechallenge, demonstrating that these mice had developed robust immunological memory against B16F10 tumor antigens (Fig. 6D). This result demonstrated that 1) elimination of Tregs is sufficient to boost the efficacy of Tx and 2) elimination of Tregs alone while maintaining the CD4^+^ effector population allows for the proper formation of long-term immune memory.

### Therapy induced *de novo* priming is necessary for therapeutic efficacy

Our data suggest that αCD4 mediates an increase in CD8^+^ T cell priming in the TdLN, which then leads to accumulation of newly primed CD8^+^ T cells in the tumor. However, an alternative explanation is that endogenous T cells already in the tumor locally proliferate and expand after αCD4 treatment. To test this hypothesis and assess if this intratumoral T cell expansion is critical to therapeutic efficacy, we treated tumor bearing mice with FTY720 concurrent with Tx + αCD4. FTY720 is a small molecule S1PR antagonist that prevents lymphocyte egress from lymphoid tissues, thus blocking any contributions from therapy-induced *de novo* priming to efficacy (*61*). FTY720 was initiated concurrently with the start of αCD4 treatment. To give sufficient time for the endogenous T cell response to develop before FTY720 initiation, treatment initiation was delayed two days (such that αCD4 and FTY720 were initiated on day 6 following tumor inoculation).

The addition of FTY720 to Tx + αCD4 abrogated therapeutic efficacy, with no complete responders and only minor tumor growth delay in this treatment cohort (Fig. 6E). Indeed, when we examined the tumor compartment via flow cytometry, addition of FTY720 to Tx + αCD4 dropped CD8^+^ T cell counts back to baseline (PBS/DMSO) levels (Fig. 6F). This confirmed that increases in CD8^+^ T cells in the tumor after Tx + αCD4 were due to *de novo* priming and trafficking from the TdLN and not local proliferation of T cells already in the tumor. The increased activation and proliferation in the TdLN (as measured by increased Ki67^+^ CD8^+^ T cells, increased CD25^+^ CD8^+^ T cells, and a shift to an effector/effector memory phenotype in the CD8^+^ T cell population) was preserved with the addition of FTY720, confirming that FTY720 prevented trafficking of these newly primed T cells to the tumor (Fig. 6G, Fig. 7A). Indeed, beginning αCD4 therapy 8 days before tumor inoculation maintained some efficacy of the combination; however, delaying initiation of αCD4 therapy to day 10 abrogated efficacy (Fig. S7B), consistent with αCD4’s role in priming (Fig. S7B).

**Figure 7.**
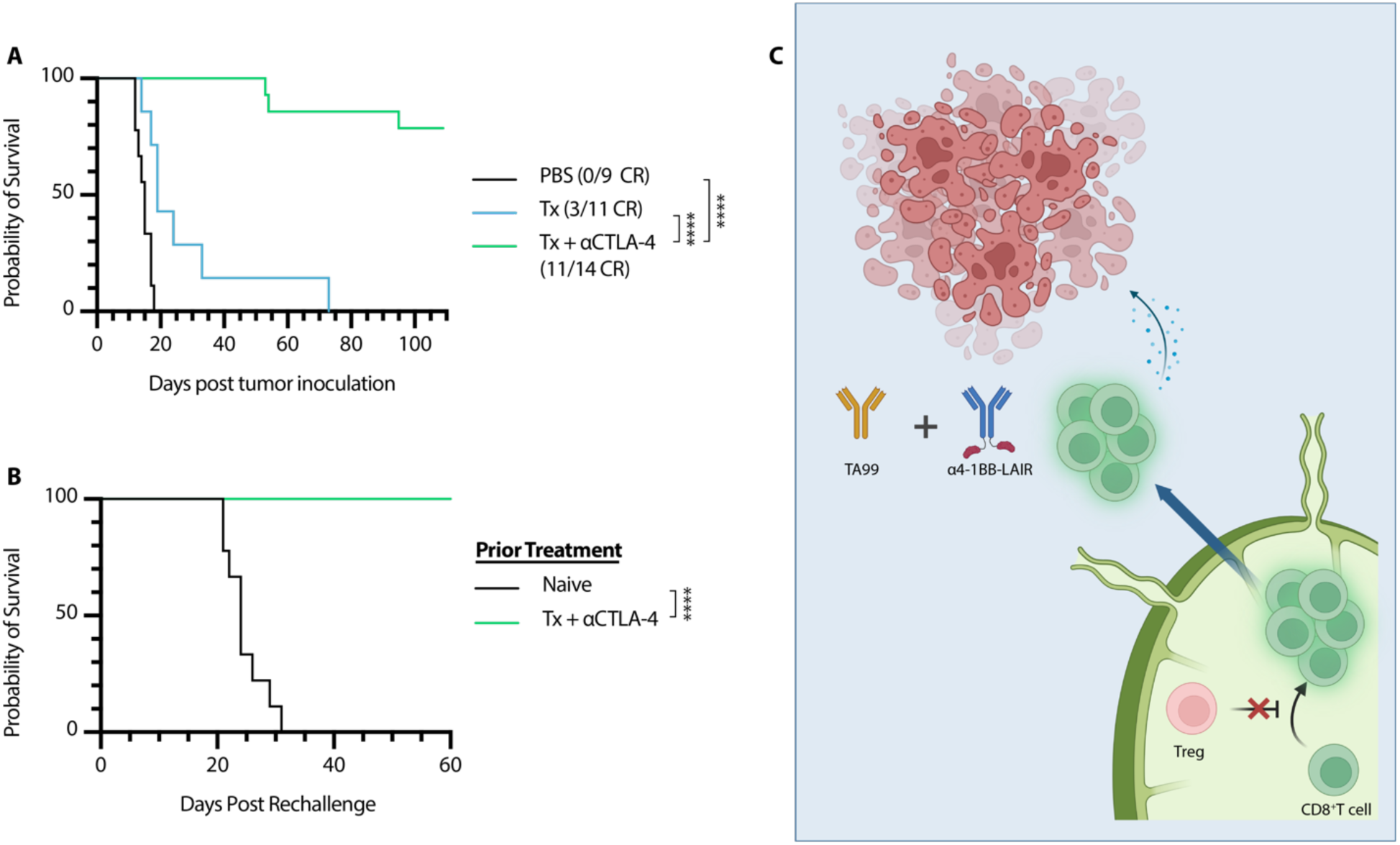
αCTLA-4 can replace αCD4 while maintaining efficacy and rescuing memory formation. Mice were inoculated with 1 x 10^6^ B16F10 cells on day 0. **(A)** Overall survival of mice treated either with PBS (n = 9, two independent studies), Tx (n = 7), or Tx + αCTLA-4 (n = 14, two independent studies). Mice were treated with the same dose/dose schedule as in Fig. 1A with 200 µg αCTLA-4 (i.p.) given on days 6, 9, 13, 16, 20, 23, and 27. **(B)** Survival of complete responders to Tx + αCTLA-4 re-challenged on the contralateral flank >100 days after primary tumor inoculation. **(C)** Graphical Abstract of proposed mechanism of action. Tregs in the TdLN constrain proper priming of tumor reactive CD8^+^ T cells, and inhibition or depletion of these cells results in a wave of newly primed CD8^+^ T cells entering the tumor, where their cytotoxic program is supported by TA99 and collagen-anchored α4-1BB-LAIR. Survival was compared using log-rank Mantel-Cox test. \**P* < 0.05, \*\**P* < 0.01, \*\*\**P* < 0.001, \*\*\*\**P* < 0.0001.

Interestingly, if initiation of FTY720 therapy was delayed just two days (concurrent with α4-1BB-LAIR), therapeutic efficacy of this combination was restored and T cell counts in the tumor were restored to the same levels as Tx + αCD4 (Fig. S7C-E). For all FTY720 dosing schemes, blood T cell levels were significantly reduced compared to untreated mice, confirming that FTY720 was functioning as expected after treatment initiation (Fig. S7F). These data suggest that only a single priming wave is sufficient for efficacy of Tx + αCD4, and this priming wave occurs in a narrow time frame of two days following αCD4 initiation.

### αCTLA-4 therapy also synergizes with TA99 + α4-1BB-LAIR

Based on the presented data, we concluded that αCD4 synergizes with Tx by initiating a wave of *de novo* priming, that these new tumor-infiltrating T cells are supported by the local α4-1BB-LAIR agonist and TA99, and that this two-step process ultimately drives therapeutic efficacy. We therefore hypothesized that other modalities capable of improving priming, such as αCTLA-4, would also synergize well with TA99 + α4-1BB-LAIR. Although the dominant mechanism of αCTLA-4 is contested, literature supports that treatment with αCTLA-4 improves T cell priming and infiltration into the tumor (*62*). We therefore treated B16F10-bearing mice with Tx + αCTLA-4, and found that this combination was also highly efficacious, with an ~80% complete response rate (Fig. 7A, Fig. S8). We hypothesized that mice cured with Tx + αCTLA-4 would also generate robust immune memory and reject rechallenge as their CD4^+^ effector T cell pool remained intact. In agreement with this hypothesis, 100% of survivors rechallenged >100 days after initial tumor inoculation rejected this secondary tumor rechallenge (Fig. 7B).

## DISCUSSION

α4-1BB agonist antibodies have demonstrated robust efficacy as both a monotherapy and in combination with other immunotherapy agents in preclinical mouse models but have so far failed in the clinic due to dose-limiting toxicities. In this work, we set out to develop α4-1BB antibodies with tumor-restricted activity via collagen anchoring. We have previously demonstrated that fusion of collagen binding proteins lumican or LAIR to extended half-life versions of IL-2 and IL-12 improves efficacy and limits toxicities when directly injected into tumors, even in relatively collagen-sparse B16F10 melanoma tumors, such as those used in this study (*27, 28, 36*).

To generate collagen anchored α4-1BB antibodies, we fused murine LAIR1 to the C-terminus of the heavy chain of an α4-1BB agonist antibody. We tested this agonist combination with a systemic antitumor antibody, TA99. We chose this combination because 1) α4-1BB agonist Urelumab is currently being tested in combination with antitumor antibodies Cetuximab, Rituximab, and Elotuzumab and 2) a wide range of other antitumor antibodies which recognize antigens expressed on tumor cells are currently in the clinic (*63*). Antitumor antibodies have been demonstrated to improve antitumor immune responses by both generating antigenic cell debris to enhance T cell priming and also by reprogramming myeloid cells in the tumor through Fc:FcγR interactions (*64*). In agreement with preliminary phase 1 data, this combination did not result in robust efficacy in our hands, with only minor growth delay and complete responses in ~5% of treated mice (*48*). However, we unexpectedly discovered that depletion of the entire CD4^+^ T cell compartment throughout the course of this combination therapy dramatically improved response rates, with >90% of mice achieving durable complete responses. A similarly efficacious combination (systemic α4-1BB + αCD4) has been reported in the literature, although durable responses were not seen, with all mice succumbing to their tumors between day 70-80 (*49*). To our knowledge, this is the highest complete response rate seen of any α4-1BB agonist antibody therapy in the poorly immunogenic B16F10 melanoma tumor model.

As Tregs comprise a sizable portion of the CD4^+^ T cell compartment, we tested Treg depletion in lieu of whole CD4^+^ T cell depletion using Foxp3-DTR mice in combination with TA99 + α4-1BB-LAIR and observed equivalent efficacy. While Tregs play a crucial role in preventing autoimmunity, they also constrain productive antitumor immune responses. Intratumoral Treg infiltration is correlated with poor prognosis across many different tumor types and there is evidence that intranodal Tregs infiltration is a better predictor of survival than blood or intratumoral Tregs in certain contexts (*65–68*). Tregs exert their effects through multiple different pathways, including secretion of immunosuppressive cytokines such as IL-10, Transforming Growth Factor-beta (TGF-β), and IL-35, acting as a sink for IL-2 due to their high expression of the IL-2 high affinity receptor CD25, generation of immunosuppressive adenosine through expression of CD39, and expression of inhibitory receptors such as CTLA-4 and LAG-3 (*43, 44*).

Tregs are a major contributor to the immunosuppressive environment of the tumor, but they can also interfere with CD8^+^ T cell priming in lymphoid tissues (*69, 70*). Even prior to the identification of the transcription factor Foxp3 as the canonical driver of Tregs, seminal work found that depletion of CD25^+^ T cells (a subset of which are Tregs) before tumor implantation can lead to enhanced antitumor immune responses and eventual spontaneous tumor rejection (*71*). Although how Tregs constrain priming is multifaceted, it is well established that CTLA-4 expressed on Tregs can transendocytose CD80 and CD86 off the surface of dendritic cells, hampering their ability to provide proper co-stimulation and prime CD8^+^ T cells (*72–74*). Blocking this transendocytosis is thought to at least partially explain the mechanism of how αCTLA-4 therapy functions to improve priming. Indeed, we show αCTLA-4 synergized as well as complete CD4^+^ T cell or Treg specific depletion with TA99 + α4-1BB-LAIR. Our work supports the notion that intranodal Tregs dampen antitumor immune responses by constraining proper priming, and that relieving this constraint can bolster the magnitude of the antitumor T cell response and synergize robustly with T cell directed agonist immunotherapies, particularly in immunologically cold tumors such as the one used in this study.

Although long-term CD4^+^ T cell compartment depletion leads to obvious defects in both T and B cell adaptive immune responses, transient CD4^+^ T cell depletion has been clinically tested in cancer and other disease states using an αCD4 antibody. Transient αCD4 depletion resulted in similar increases in *de novo* priming and CD8^+^ T cell infiltration in the tumor, consistent with our own data (*52, 53*). However, although no adverse events have been reported in these small phase 1 trials, these patients are still at risk of severe and possibly fatal infections if exposed to pathogens while devoid of their CD4^+^ compartment. Additionally, although αCD4 depletion therapy synergized well with TA99 + α4-1BB-LAIR, mice failed to form immunological memory, which can be important for long term tumor control and control of distant metastases. In patients, the presence of memory T cells corresponds with breast cancer survival and memory T cells have been reported to persist in survivors of melanoma treated with immunotherapy for at least 9 years (*75, 76*). With this in mind, we set out to understand the mechanism of how CD4^+^ T cell compartment depletion synergized with TA99 + α4-1BB-LAIR and develop new combination therapies with higher translational potential. Our data demonstrated that CD4^+^ T cell depletion eliminated Tregs in the TdLN, removing immunosuppressive constraints on proper CD8^+^ T cell priming, and induced a wave of freshly primed T cells to enter the TME. The combination of TA99 + α4-1BB-LAIR is able to reprogram the TME into a more supportive environment for these newly primed T cells, allowing them to maintain their cytotoxic phenotype, leading to tumor regression and clearance (Fig. 7C). Indeed, recent data has suggested a two-step model for CD8^+^ T cell activation in cancer, with initial activation in the TdLN and effector differentiation occurring with co-stimulation in the tumor (*77*). The two components of our therapy mirror this paradigm, with αCD4 increasing activation in the TdLN and TA99 + α4-1BB-LAIR enhancing effector functions of these newly activated CD8^+^ T cells directly in the tumor.

This localized therapy is reliant on intratumoral administration of the α4-1BB-LAIR component, which is clinically feasible with advances in interventional radiology (*78–80*). Indeed, the oncolytic virus therapy talimogene laherparepvec (T-vec) has been approved since 2015 and is routinely injected into cutaneous and subcutaneous unresectable melanoma lesions (*81, 82*). Preclinical and clinical development around intratumorally administered therapies have been steadily on the rise.

This study has the potential for immediate translational impact. Since both antitumor antibodies and αCTLA-4 antagonists are approved and routinely used in clinically, they could easily be combined with clinical stage localized α4-1BB agonists. Indeed, our data demonstrated that even non-collagen anchored α4-1BB agonists synergize fairly well with antitumor antibodies in combination with αCTLA-4 therapy, identifying a potential triple combination therapy whose individual components are all already in clinical use.

In conclusion, we found that effective TA99 + α4-1BB-LAIR therapy requires a wave of *de novo* CD8^+^ T cell priming to achieve maximum efficacy. In this study, we generated this enhanced priming wave through whole CD4^+^ T cell compartment depletion with an αCD4 depleting antibody, Treg specific ablation using Foxp3-DTR mice, or treatment with αCTLA-4, a modality known to increase priming. These combinations resulted in high levels of primary tumor efficacy, with ~80-100% complete response rates. However, only in the latter two strategies, which preserved CD4^+^ effector T cells, did mice also develop robust long-term immunological memory, with 100% of cured mice rejecting secondary tumor rechallenge. Our data demonstrate that at baseline, proper CD8^+^ T cell priming is constrained by Tregs present in the TdLN. All three priming enhancing strategies are directed towards Tregs, either depleting them completely (αCD4 and DT), or blocking their immunosuppressive pathways (αCTLA-4). This provides strong rationale for development of Treg-directed therapies that modulate Treg function in the TdLN which, in combination with proper immune agonists, can drive high levels of efficacy even in immunologically cold tumors.

## MATERIALS AND METHODS

### Study Design

The purpose of this study was to (i) evaluate the efficacy and safety of collagen anchoring α4-1BB-LAIR and subsequently to (ii) understand the mechanism driving synergy between TA99 + α4-1BB-LAIR and αCD4 and finally to (iii) identity more clinically relevant therapies that synergize with TA99 + α4-1BB-LAIR. We used the syngeneic murine melanoma line B16F10 for all studies. Mice were randomized before beginning treatment to ensure equal tumor size in all groups and were monitored for tumor size and weight loss until euthanasia or until complete tumor regression. Investigators were not blinded during the studies. In all studies there were at least 5 mice per experimental group, except for the bulk RNA-sequencing experiment which had 3-4 mice per group. No data/experiments were excluded unless there were technical issues with the experiment, and outliers were not excluded. Many experiments were repeated twice, and number of mice per group, number of experimental repeats, and statistical methods are noted in figure legends.

### Mice

C57Bl/6 (C57Bl/6NTac) mice were purchased from Taconic. C57Bl/6 albino (B6(Cg)-Tyr^c-2J^/J) mice were purchased from The Jackson Laboratory. C57Bl/6 Foxp3-DTR(B6.129(Cg)-Foxp3^tm3(DTR/GFP)Ayr^/J) mice were a gift from the Spranger lab (MIT). B6 Foxp3-DTR mice were bred in house and genotyped using Transnetyx. All animal work was conducted under the approval of the Massachusetts Institute of Technology Committee on Animal Care in accordance with federal, state, and local guidelines.

### Cells

B16F10 cells were purchased from ATCC. Apigmented B16F10 cells used for imaging were generated by genetic deletion of Tyrosinase-related-protein-2 (TRP2), referred to as B16F10-Trp2 KO cells (*83*). Tumor cells were cultured in Dulbecco’s Modified Eagle Medium (DMEM, ATCC) supplemented with 10% Fetal Bovine Serum (FBS, Gibco). FreeStyle 293-F cells and Expi293 cells were purchased from Invitrogen and cultured in FreeStyle expression medium (Gibco) and Expi293 expression medium (Gibco), respectively. CHO DG44 cells were cultured in ProCHO5 (Lonza) supplemented with 4 mM L-glutamine, 0.1 mM hypoxanthine, and 16 μM thymidine. Tumor cells were maintained at 37°C and 5% CO_2_ and FreeStyle 293-F cells, Expi293 cells, and CHO DG44 cells were maintained at 37°C and 8% CO_2_. All cells tested negative for mycoplasma contamination.

### Tumor Inoculation and Treatment

Mice were aged six to twelve weeks before tumor inoculations. 1 x 10^6^ B16F10 or B16F10-Trp2KO cells were suspended in 50uL sterile PBS (Corning) and injected subcutaneously on the right flank.

Mice were randomized before beginning treatment to ensure equal tumor size in all groups. TA99 was administered intraperitoneally (i.p) at a dose of 200 μg in 200 μL sterile PBS (Corning). α4-1BB or α4-1BB-LAIR was administered intratumorally (i.t.) in 20 μL of sterile PBS (Corning) at a dose of 30 μg or 36.1 μg (molar equivalents), respectively. αCD4 (Clone GK1.5, Bioxcell) was administered i.p. at a dose of 400 μg in 100 μL sterile PBS (Corning). αCTLA-4 (Clone 9D9, mIgG2c isotype) was administered i.p. at a dose of 200 μg in 100 μL of sterile PBS (Corning). Diphtheria Toxin (DT, Sigma Aldrich) was administered i.p. at a dose of 1 μg in 100 μL sterile PBS (Corning) or i.t. at a dose of 75 ng or 125 ng in 20 μL sterile PBS (Corning). Stock solutions of FTY720 (Sigma Aldrich) were resuspended at 10 mg/mL in DMSO and diluted to a dose of 30 μg in sterile PBS (Corning) to a final volume of 150 μL and administered i.p.

TA99 was dosed on days 5, 12, 19, and 26 and α4-1BB and α4-1BB-LAIR were administered on days 6, 13, 20, and 27. αCD4 was administered starting on day 4 and continued every three days until day 37 (Fig. 1A). For some studies, therapy initiation was delayed by 2 days to allow for larger tumors at time of analysis (flow cytometry, chemokine/cytokine analysis, and bulk-RNA-sequencing), sufficiently sized tumors for intratumoral DT administration (DT survival studies), or to avoid interfering with the endogenous T cell response (FTY720 studies, except Fig. S7C which followed Fig. 1A dosing scheme). DT was administered every other day starting on day 6 and continued until day 36. FTY720 was administered starting concurrently with αCD4 and continued every other day until one week after final α4-1BB-LAIR dose. “Delayed” FTY720 was administered starting concurrently with α4-1BB-LAIR and continued every other day until one week after final α4-1BB-LAIR dose. αCTLA-4 was given on days 6, 9, 13, 16, 20, 23, and 27.

During all tumor studies, mice were monitored continuously for tumor growth and weight change. Tumor growth was assessed by direct measurement with calipers and mice were euthanized when their tumor area (length × width) reached 100 mm^2^ or mice lost more than 20% of their body weight. Mice that were cured of their primary tumor but later euthanized due to overgrooming related dermatitis were still classified as complete responders and included in analysis.

For rechallenge studies, mice that rejected their primary tumors were inoculated with 1 x 10^5^ B16F10 tumor cells on the left, or contralateral, flank 100-110 days after primary tumor inoculation and monitored for tumor outgrowth. Age matched naïve mice were used as controls in these studies

### Flow Cytometry

Tumors were excised, weighed, and mechanically dissociated before being enzymatically digested using a gentleMACS Octo Dissociator with Heaters (Miltenyi Biotec) in gentleMACS C tubes (Miltenyi Biotec) and enzymes from the Mouse Tumor Dissociation Kit (Miltenyi Biotec). Tumors were digested using the 37C_m_TDK_1 program for soft tumors. Following digestion, tumors were filtered through a 40 µm filter and transferred to a V-bottom 96 well plate for staining. TdLN and spleens were excised, weighed, and mechanically dissociated through a 70 µm filter. Spleen samples were resuspended with 5 mL of ACK Lysis buffer (Gibco) to lyse red blood cells before being re-filtered through a 70 µm filter. TdLN and spleen samples were then transferred to a V-bottom 96 well plate for staining. Blood samples were collected via cardiac puncture into K3 EDTA coated tubes (MiniCollect). 200 µL of blood was mixed with 1 mL of ACK lysis buffer (Gibco) to lyse red blood cells before being transferred to a V-bottom 96 well plate for staining. Precision Counting Beads (Biolegend) were added to each well to account for sample loss during processing and obtain accurate counts. Cells were washed once with PBS and then resuspended in Zombie UV Fixable Viability Dye (Biolegend) to stain dead cells for 30 minutes at RT in the dark. Cells were then washed with FACS buffer (PBS (Corning) + 0.1% BSA (Sigma Aldrich) + 2mM EDTA (Gibco)) and blocked with αCD16/CD32 antibody (Clone 93, eBioscience) for 20 minutes on ice in the dark and then stained for extracellular markers for 30 minutes on ice in the dark. Samples not requiring intracellular staining were washed with FACS buffer and fixed with BD Cytofix (BD Biosciences) for 30 minutes at RT in the dark. Cells were then washed and resuspended in FACS buffer. For samples requiring intracellular staining, cells were washed after extracellular staining, fixed and permeabilized with the Foxp3/Transcription Factor Staining Buffer Set (eBiosciences), and stained for 30 minutes at RT in the dark, before being washed and resuspended in FACS buffer. Samples were analyzed with a BD FACS Symphony A3 (BD Biosciences) and data was processed and analyzed with FlowJo V10. See Figure S9 for example gates.

Tumor and TdLN samples in Figures 2 and S4 were stained with αCD45-BUV395 (30-F11, BD Biosciences), αCD4-BUV563 (RM4-4, BD Biosciences), αCD8α-BUV737 (53-6.7 BD Biosciences), αCD62L-BUV805 (MEL-14, BD Biosciences), αCD44-BV421 (1M7, Biolegend), αKi67-BV605 (16A8, Biolegend), αCD3-BV711 (17A2, Biolegend), αTIM-3-BV785 (RMT3-23, Biolegend), αTCF1/TCF7-AF488 (C63D9, Cell Signaling Technology), αPD-1-PerCp/Cy5.5 (29F.1A12, Biolegend), αFoxp3-PE (FJK-16s, Invitrogen), αCD25-PE-Cy5 (PC61, Biolegend), αNK1.1-PE-Cy7 (PK126, Biolegend), α4-1BB-APC (17B5, Biolegend), αCD107a-APC-Cy7 (1D4B, Biolegend).

Tumor, TdLN, and spleen samples in Figure 5A and S6 were stained with αCD45-BUV395 (30F-11, BD Bioscience), αCD8a-BUV737 (53-6.7, BD Biosciences), αCD3-BV785 (17A2, Biolegend), αNK1.1-PE-Cy7 (PK-136, Biolegend), αCD4-APC-Cy7 (GK1.5, Biolegend), and Foxp3^+^ cells were identified using the GFP reporter expressed under the *Foxp3* locus in Foxp3-DTR mice.

Tumor, TdLN and blood samples in Figures 5F-G and S7 were stained with αCD45-BUV395 (30-F11, BD Biosciences), αCD4-BUV563 (RM4-4, BD Biosciences), αCD44-BUV737 (1M7 BD Biosciences), αKi67-BV421 (16A8, Biolegend), αCD3-BV711 (17A2, Biolegend), αCD8a-FITC (53-6.7, Biolegend) αFoxp3-PE (FJK-16s, Invitrogen), αCD25-PE-Cy5 (PC61, Biolegend), αNK1.1-PE-Cy7 (PK126, Biolegend), αCD62L-APC (MEL-14, Biolegend), αCD107a-APC-Cy7 (1D4B, Biolegend).

### RNA extraction for Sequencing

Tumor samples were processed as previously described. Samples were enriched for CD45^+^ cells using an EasySep Mouse TIL (CD45) Positive Selection kit (STEMCELL) and RNA was extracted with an RNeasy Plus Mini Kit (Qiagen). TdLN samples were processed as previously described. Samples were again enriched for CD45^+^ cells using an EasySep Mouse CD45 Positive Selection kit (STEMCELL) and RNA was extracted with an RNeasy Plus Mini Kit (Qiagen). RNA was stored at −80°C until further processing.

### RNA-seq Library Preparation and Sequencing

RNA-sequencing was performed by the BioMicro Center at MIT using a modified version of the SCRB-seq protocol (*84*). Libraries were sequenced on a NextSeq 500 using a 75-cycle kit.

### RNA-seq Alignment, Quantification, and Quality Control

Data preprocessing and count matrix construction were performed using the Smart-seq2 Multi-Sample v2.2.0 Pipeline (RRID:SCR_018920) on Terra. For each cell in the batch, single-end FASTQ files were first processed with the Smart-seq2 Single Sample v5.1.1 Pipeline (RRID:SCR_021228). Reads were aligned to the GENCODE mouse (M21) reference genome using HISAT2 v2.1.0 with default parameters in addition to --k 10 options. Metrics were collected and duplicate reads marked using the Picard v.2.10.10 CollectMultipleMetrics and CollectRnaSeqMetrics, and MarkDuplicates functions with validation_stringency=silent. For transcriptome quantification, reads were aligned to the GENCODE transcriptome using HISAT2 v2.1.0 with --k 10 --no-mixed --no-softclip --no-discordant --rdg 99999999,99999999 --rfg 99999999,99999999 --no-spliced-alignment options. Gene expression was calculated using RSEM v1.3.0’s rsem-calculate-expression --calc-pme --single-cell-prior. QC metrics, RSEM TPMs and RSEM estimated counts were exported to a single Loom file for each sample. All individual Loom files for the entire batch were aggregated into a single Loom file for downstream processing. The final output included the unfiltered Loom and the tagged, unfiltered individual BAM files.

### RNA-Seq Analysis

Samples with less than 10,000 genes detected were excluded from analysis. This led to exclusion of two tumor samples, one from the Tx + αCD4 group and one from the αCD4 group at the day 6 time point (Fig. S10). UMAP embedding of TdLN samples was generated from the top 5 principal components and top 3000 variable features. *DEseq2* was used to conduct differential expression testing and *apeglm* was used for effect size estimation (85, 86). Pathways enrichment analysis for statistically significant upregulated genes was performed using *enrichR* to query the databases indicated in the text (*54–56*). A score for the derived response gene signature was calculated for each experimental cohort using Seurat (*AddModuleScore*) (*87*). Differential expression testing was performed as described above comparing all tumor sample cohorts to the D3 PBS, D6 PBS, D3 Tx + aCD4, and D6 Tx + aCD4 cohorts. All statistically significant hits (*p*-adj ≦5 with absolute value log2 fold-change ≥ 2 were included for further analysis. Gene clusters were defined using *k*-means clustering and the *complexHeatmap* package was used to generate expression heatmaps for these genes (*88*). Relative expression profiles of these gene clusters were generated by summarizing the percent expression using Seurat (*PercentageFeatureSet*) per sample and dividing by the highest average percent per condition (*87*). Gene sets were obtained from MSigDB and enrichment of genes from each cluster in these gene sets was calculated using the *enrichGO* package (*89*).

### Statistical Methods

Statistics were computed in GraphPad Prism v9 as indicated in figure captions. Survival studies were compared using the log-rank (Mantel-Cox) test. Flow data and tumor supernatant cytokine/chemokine data were compared using one- or two-way ANOVA with Tukey’s multiple comparison correction. Differential expression analysis using DESeq2 models counts for each gene using a negative binomial model and tests for significance using Wald tests (*86*). Gene set enrichment is calculated by the Fisher’s exact test. *P* values are corrected for multiple hypothesis testing using the Benjamini-Hochberg procedure for all RNA-sequencing analysis. Sample size and *P*-value cutoffs are indicated in figure captions.

## Supplementary Materials

Materials and Methods

Figs. S1 to S10

Table S1

Data files S1 to S8

## Acknowledgement

We thank the Koch Institute’s Robert A. Swanson (1969) Biotechnology Center (National Cancer Institute Grant P30-CA14051) for technical support, specifically the Flow Cytometry Core Facility and the BioMicro Center. We thank the Spranger lab for the gift of the Foxp3-DTR breeding pair. We thank the Protein Production and Structure Core Facility at the École polytechnique fédérale de Lausanne for the development of the 9D9 stable line. Figures partially created with biorender.com.

## Author Contributions

Conceptualization: JRP, KDW

Designed Experiments: JRP, KDW

Conducted Experiments: JRP, BML, LD, JAS, LS, EAL, WP

Analyzed Data: JRP, BML, JMP

Wrote manuscript: JRP, KDW

## Data and materials availability

All raw data or materials are available upon request. Sequencing data can be found in the GEO database under accession GSE223087.

## SI Methods

### Cloning and Protein Production

The heavy chain and light chain variable regions of α4-1BB antibody (clone LOB12.3) were synthesized as gBlock gene fragments (Integrated DNA technologies) and cloned into the gWiz expression vector (Genlantis) using In-fusion cloning (Takara Bio). Antibodies were expressed as chimeras with a murine kappa light chain constant region and a murine IgG1 heavy chain constant region. Antibodies were encoded in a single expression cassette with a T2A peptide inserted between the light chain and heavy chain. αFITC (clone 4420) were constructed in the same fashion, but a murine IgG2c isotype with LALA-PG silencing mutations was used for the heavy chain constant region (90). For LAIR fusions, the murine LAIR1 gene was synthesized as a gBlock gene fragment (Integrated DNA technologies) and cloned as a fusion to the C-terminus of the heavy chain constant region separated by a flexible (G_4_S)_3_ linker. Plasmids were transformed into Stellar competent cells (Takara Bio) for amplification and isolated with Nucleobond Xtra endotoxin-free kits (Macherey-Nagel).

a4-1BB, a4-1BB-LAIR, αFITC, and αFITC-LAIR were produced using the Expi293 expression system (Gibco) following manufacturer’s instructions. Briefly, 1 mg/L of DNA and 3.2 mg/L of ExpiFectamine 293 were individually diluted into OptiMEM media (Gibco) and then combined dropwise. This mixture was then added dropwise to Expi293F suspension cells and 18-24 hours later ExpiFectamine 293 Transfection enhancers 1 and 2 (Gibco) were added to the culture. 7 days after transfection supernatants were harvested and antibodies were purified using Protein G sepharose 4 Fast Flow resin (Cytiva).

TA99 was produced using a FreeStyle 293-F stable production line generated in-house. Cells were expanded and then seeded at a density of 1 M/mL and supernatant was harvested 7 days later. 9D9 was produced using a CHO DG44 stable production line gifted to us by David Hacker. Cells were expanded and then seeded at a density of 0.5M/mL and supernatant was harvested 7 days later. Both TA99 and 9D9 were purified using rProtein A Sepharose Fast Flow resin (Cytiva).

Following purification, proteins were buffer exchanged into PBS (Corning) using Amicon Spin Filters (Sigma Aldrich), 0.22 µm sterile filtered (Pall), and confirmed for minimal endotoxin (<0.1 EU/dose) using the Endosafe LAL Cartridge Technology (Charles River). Molecular weight was confirmed with SDS-PAGE. Proteins were run alongside a Novex Sharp Pre-Stained Protein Standard (Invitrogen) on a NuPAGE 4 to 12% Bis-Tris gel (Invitrogen) with 2-(*N*-morpholino) ethanesulfonic acid (MES) running buffer (VWR) and stained for visualization with SimplyBlue Safe Stain (Life Technologies). Proteins were confirmed to be free of aggregates by size exclusion chromatography using a Superdex 200 Increase 10/300 GL column on an Äkta Explorer FPLC system (Cytiva). All proteins were flash frozen in liquid nitrogen and stored at −80°C.

### IVIS

Proteins were labeled with Alexa Fluor 647 NHS Ester (Life Technologies) and a Zeba desalting column (Thermo Scientific) was used to remove excess dye. Total molar amount of dye injected per sample was normalized between groups before injection. 20 µg of αFITC mIgG2c LALA-PG and a molar equivalent of αFITC-LAIR mIgG2c LALA-PG were used for *in vivo* retention studies. C57Bl/6 albino mice were inoculated with 10^6^ B16F10-Trp2 KO cells and labeled proteins were injected i.t. on day 7. Fluorescence at the site of the tumor was measured longitudinally using the IVIS Spectrum Imaging System (Perkin Elmer). One week prior to study initiation, mice were switched to an alfalfa-free casein chow (Test Diet) to reduce background fluorescence. Total radiant efficiency was calculated after subtracting background fluorescence and normalizing to the maximum value for each protein using Living Image software (Caliper Life Sciences).

### Tumor Cytokine/Chemokine Analysis

Tumors were excised, weighed, mechanically dissociated, and incubated in tissue protein extraction reagent (T-PER, Thermo Fisher Scientific) with 1% Halt protease and phosphatase inhibitors (Thermo Fisher Scientific) for 30 minutes at 4°C while rotating. The lysates were then centrifuged and supernatants filtered through a Costar 0.22 µm SpinX filter (Corning) to remove any remaining debris. Lysates were flash frozen and stored at −20°C until time of analysis. Lysates were analyzed with the 13-plex mouse Cytokine Release Syndrome LEGENDplex panel and the Mouse/Rat Total/Active TGF-β1 LEGENDplex kit (Biolegend). Data was collected on a BD LSR II cytometer (BD Biosciences).

### Collagen I ELISA

96 well plates precoated with rat collagen I (Gibco) were blocked overnight with PBSTA (PBS (Corning) + 2% w/v BSA (Sigma Alrich) + 0.05% v/v Tween-20 (Millipore Sigma)) at 4°C. After washing with 3 times PBST (PBS (Corning) + 0.05% v/v Tween-20 (Millipore Sigma)) and 3 times with PBS (Corning), a4-1BB and a4-1BB-LAIR were incubated in PBSTA overnight at 4°C while shaking. Wells were washed 3 times with PBST and 3 times with PBS and then incubated with goat αmIgG1-Horseradish peroxidase (HRP) (1:2000, Abcam) in PBSTA for 1 hour at RT while shaking. Wells were again washed 3 times with PBST and 3 times with PBS and then 1-Step Ultra TMB-ELISA Substrate Solution (Thermo Fisher) was added for 5-15 min, followed by 1 M sulfuric acid to quench the reaction. Absorbance at 450 nm (using absorbance at 570 nm as a reference) was measured on an Infinite M200 microplate reader (Tecan). Binding curves were generated with GraphPad Prism software V9. K_D_ values were calculated using a nonlinear regression fit for one site total binding with no non-specificity and curves were normalized to the B_max_ values.

### Surface 4-1BB Binding Assay

The gene for murine 4-1BB (OriGene) was cloned into the pIRES2 expression vector, which encodes for GFP downstream of the inserted 4-1BB gene using an IRES site, using In-Fusion cloning (Takara Bio). Freestyle 293-F cells were transiently transfected by mixing 1 mg/mL of plasmid DNA and 2 mg/mL of polyethylenimine (Polysciences) in OptiPRO Serum Free Medium (Gibco) and, after incubating, adding dropwise to the cells. 3-5 days after transfection, cells were harvested and pelleted in V-bottom 96 well plates. Cells were titrated with a4-1BB or a4-1BB-LAIR and incubated for 3 hours shaking at 4°C. Cells were washed with PBSA (PBS (Corning) + 0.1% BSA (Sigma Aldrich)) and incubated with αmIgG1-APC (diluted 1:250, clone M1-14D12, Biolegend) for 30 minutes shaking at 4°C. Data was collected on a BD LSR II cytometer (BD Biosciences). Binding curves were generated with GraphPad Prism software V9. K_D_ values were calculated using a nonlinear regression fit for one site total binding with no non-specificity and curves were normalized to the B_max_ values.

---

**Figure S1.**
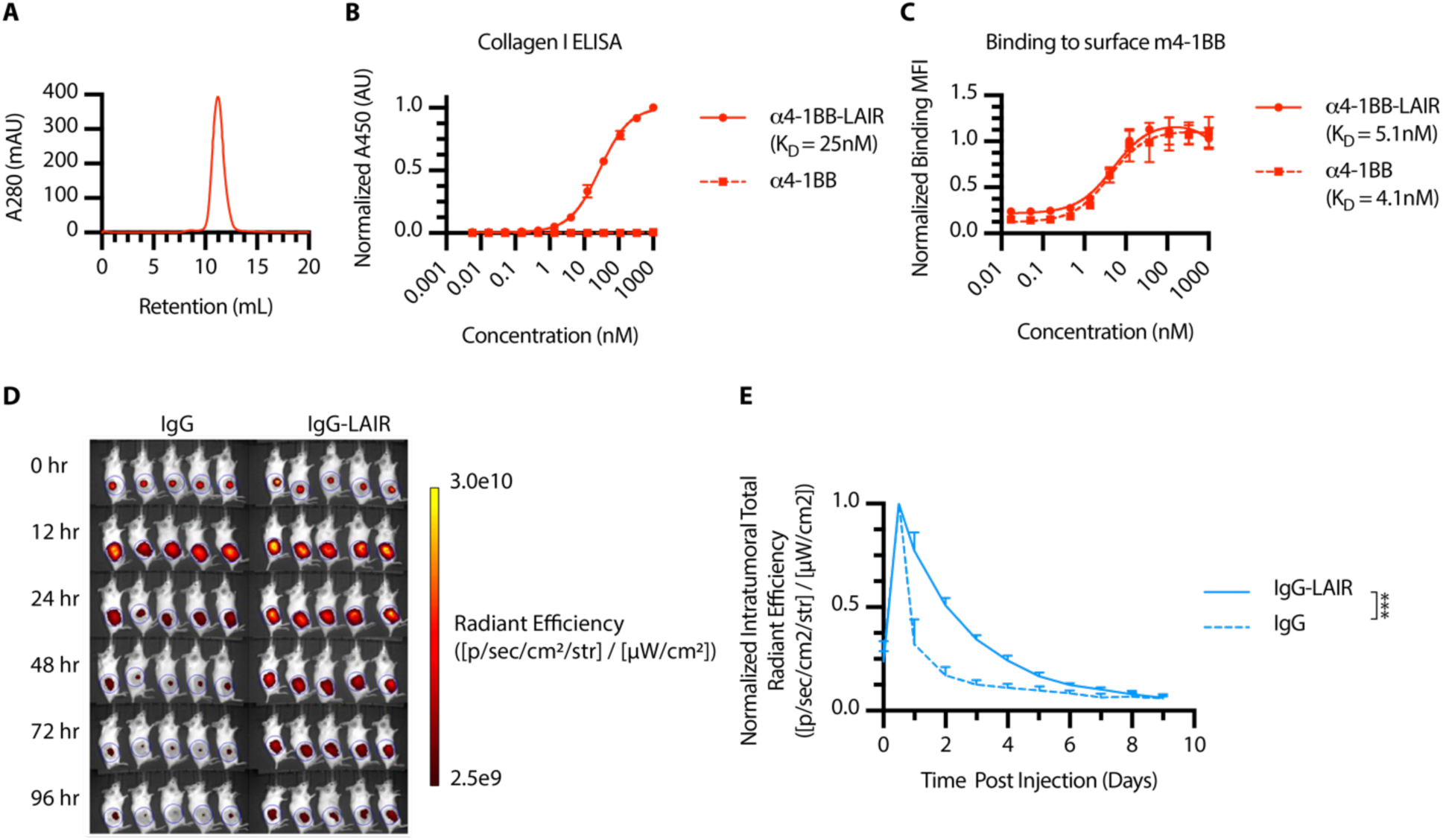
α4-1BB-LAIR behaves as expected *in vitro* and *in vivo*. **(A)** SEC chromatogram of α4-1BB-LAIR. **(B)** Equilibrium binding curve of α4-1BB-LAIR and α4-1BB to collagen I coated plates (mean ± S.D., n = 4). **(C)** Equilibrium binding curve of α4-1BB-LAIR and α4-1BB to HEK cells expressing murine 4-1BB (mean ± S.D., n = 4). **(D-E)** Mice were inoculated with 1 x 10^6^ B16F10-Trp2 KO cells on day 0, injected with 20 µg of fluorescently labeled control IgG or equimolar amount of IgG-LAIR, and fluorescence was measured longitudinally via IVIS. **(D)** example fluorescence images from select timepoints and **(E)** Quantification of normalized radiant efficiency (mean ± S.D.) in mice receiving IgG or IgG-LAIR (n = 5). Retention data were compared using two way ANOVA with Tukey’s multiple hypothesis testing correction. \**P* < 0.05, \*\**P* < 0.01, \*\*\**P* < 0.001, \*\*\*\**P* < 0.0001.

**Figure S2.**
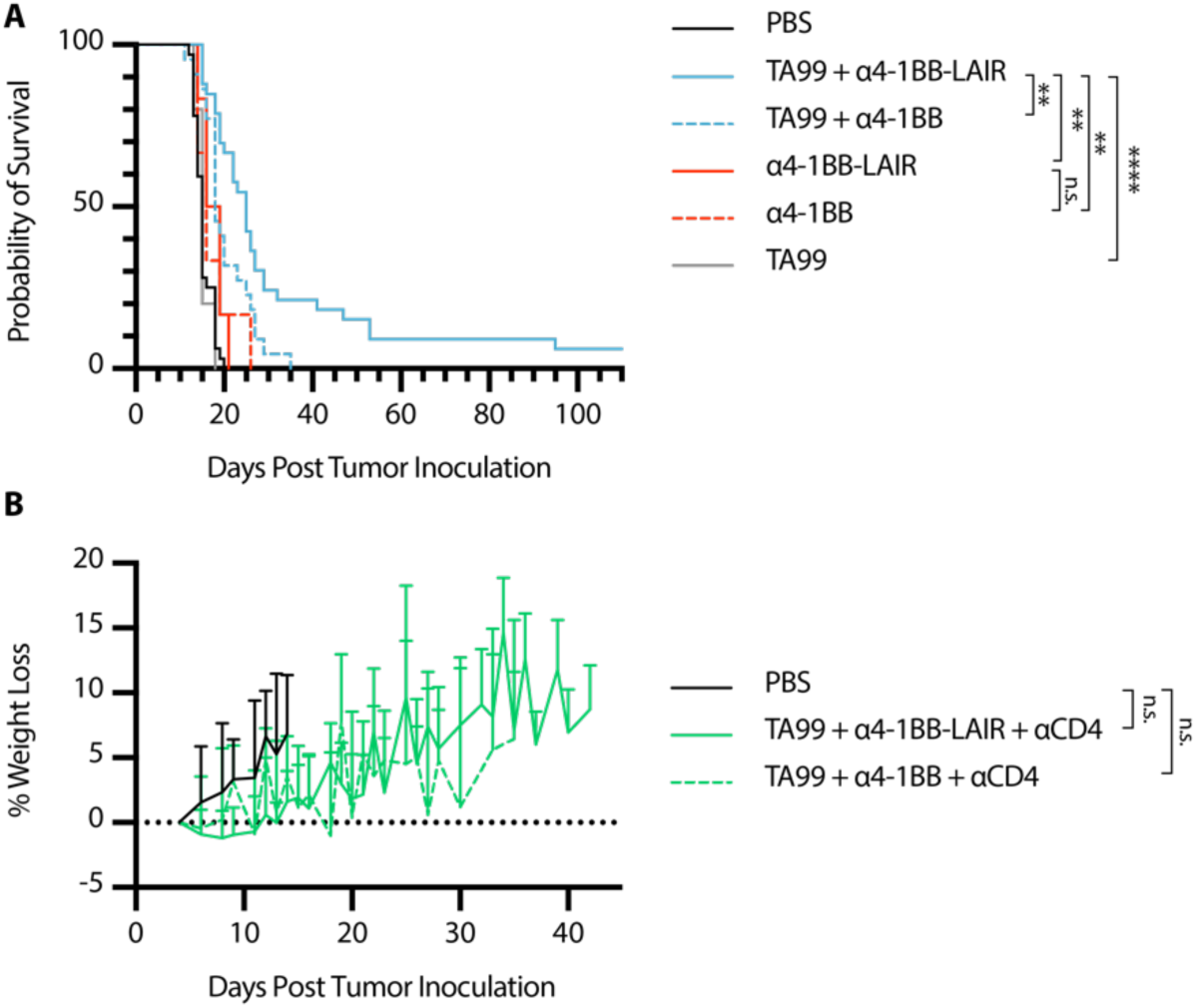
Monotherapies are not efficacious and Tx + αCD4 exhibits no toxicity. **(A)** Overall survival of mice treated with PBS (n = 32), TA99 + α4-1BB-LAIR (“Tx”, n = 33), TA99 + α4-1BB (n = 22), TA99 (n = 5), α4-1BB (n = 6), or α4-1BB-LAIR (n = 6) with treatment schedule outlined in Fig. 1A (six independent studies). **(B)** Weight loss of mice treated with PBS (n = 10), TA99 + α4-1BB-LAIR + αCD4 (n = 10), or TA99 + α4-1BB + αCD4 (n = 9) from survival study shown in Fig. 1D (two independent studies). Survival was compared using the log-rank Mantel-Cox test and weight loss data were compared using two-way ANOVA with Tukey’s multiple hypothesis testing correction. \**P* < 0.05, \*\**P* < 0.01, \*\*\**P* < 0.001, \*\*\*\**P* < 0.0001.

**Figure S3.**
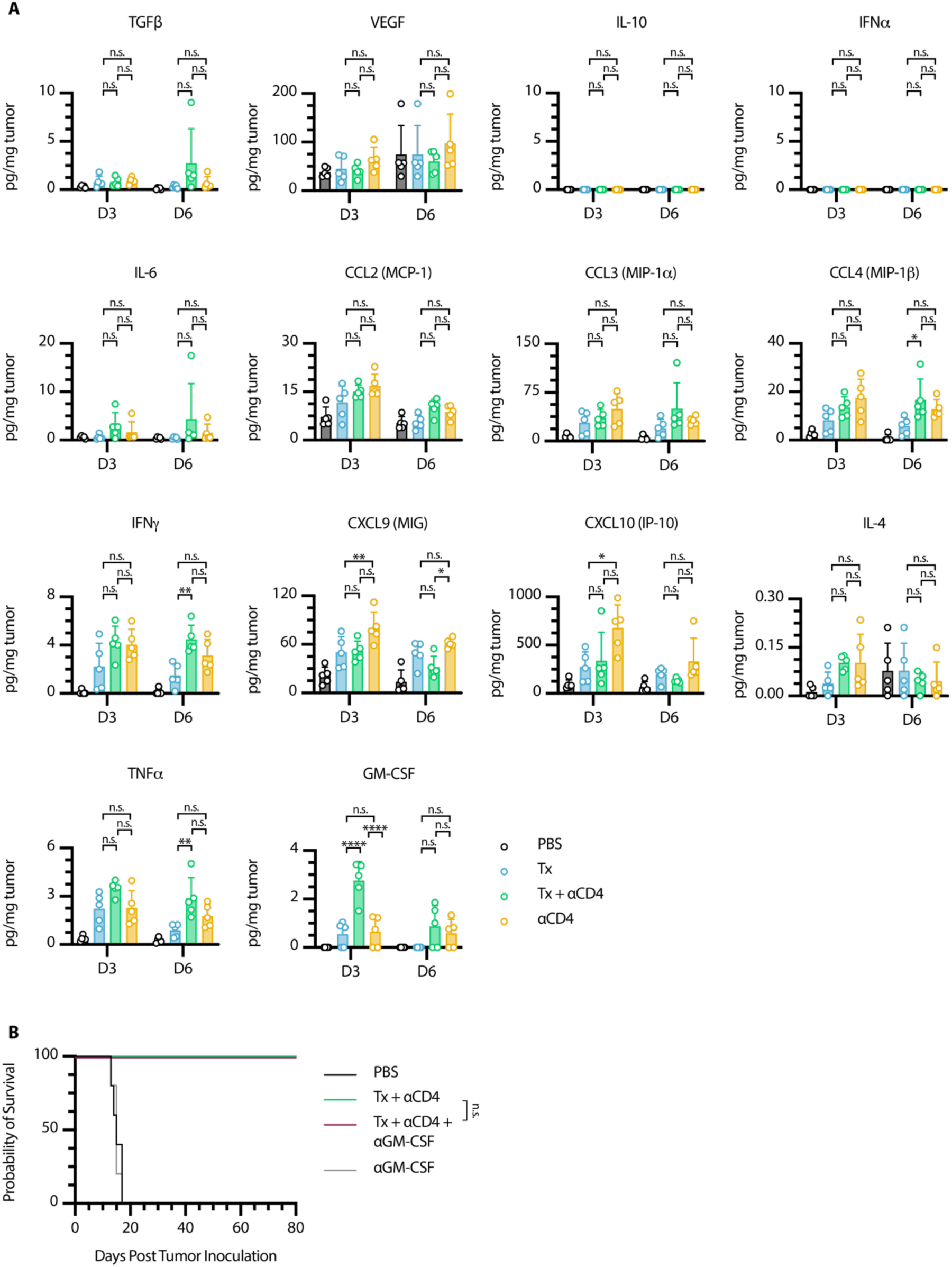
Tumor supernatant cytokine/chemokine analysis does not explain differences in efficacy. **(A)** Measured levels of indicated soluble cytokines/chemokines in tumor supernatant 3 and 6 days after first α4-1BB-LAIR treatment (n = 5). **(B)** Survival of mice treated with PBS (n = 5), Tx + αCD4 (n = 7), Tx + αCD4 + αGM-CSF (n = 7), or αGM-CSF (n = 5). Chemokine/cytokine measurements were compared using two-0way ANOVA with Tukey’s multiple hypothesis testing correction. Survival was compared using the log-rank Mantel-Cox test. \**P* < 0.05, \*\**P* < 0.01, \*\*\**P* < 0.001, \*\*\*\**P* < 0.0001.

**Figure S4.**
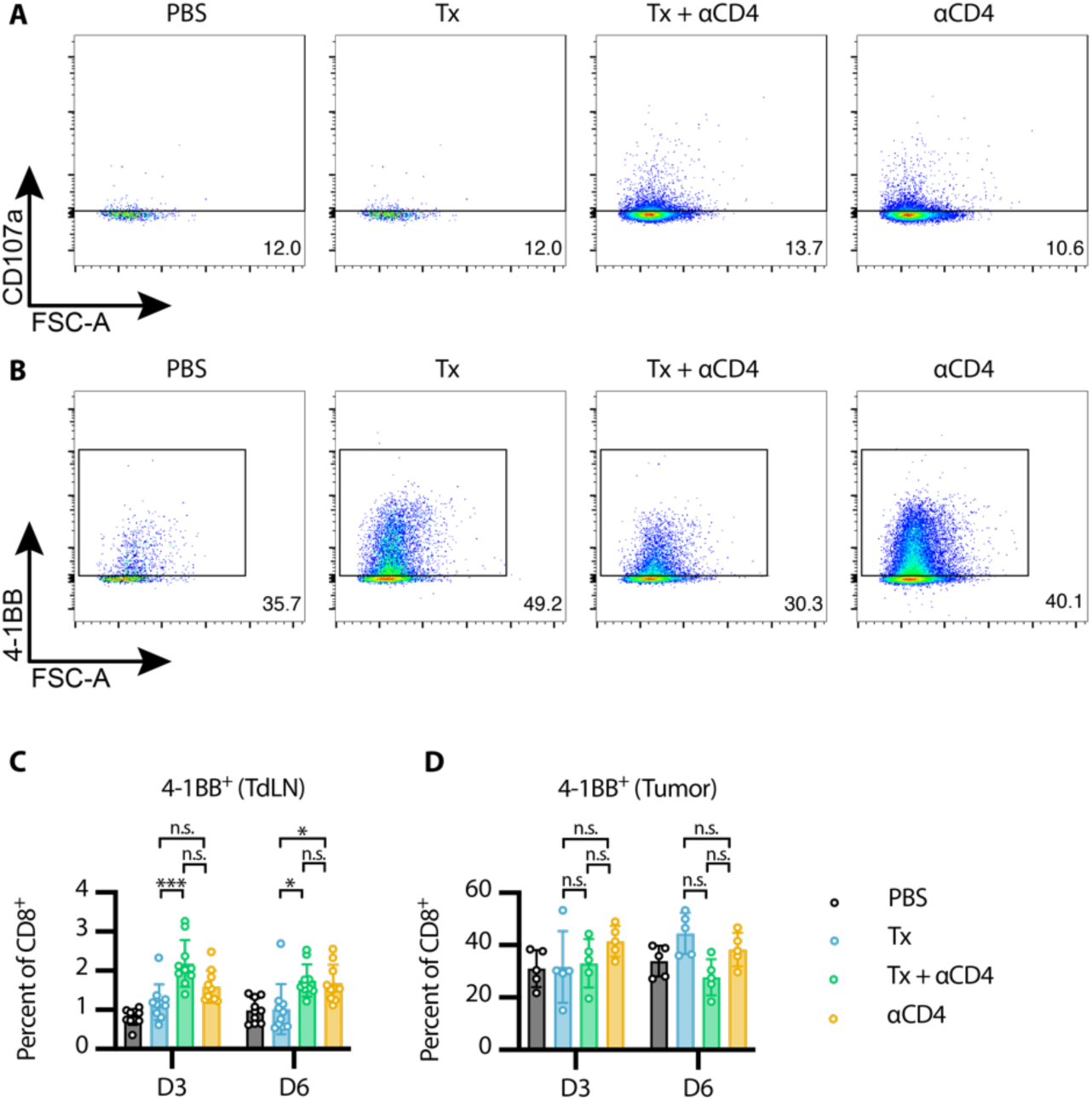
4-1BB expression on CD8^+^ TILs uniform across treatment groups. **(A)** Representative gating of CD107a^+^ CD8^+^ T cells in the tumor 6 days after first α4-1BB-LAIR treatment (gated on single cell/live/CD45^+^/CD3^+^NK1.1^−^/CD8^+^). **(B)** Representative gating of 4-1BB^+^ CD8^+^ T cells in the tumor 6 days after first α4-1BB-LAIR treatment (gated on single cell/live/CD45^+^/CD3^+^NK1.1^−^/CD8^+^). Flow cytometry quantification (mean±SD) of 4-1BB^+^ CD8^+^ T cells in the **(C)** TdLN and **(D)** tumor 3 and 6 days after first α4-1BB-LAIR treatment (gated on single cell/live/CD45^+^/CD3^+^NK1.1^−^/CD8^+^, n = 5-10, two independent experiments). Flow data were compared using two-way ANOVA with Tukey’s multiple hypothesis testing correction. \**P* < 0.05, \*\**P* < 0.01, \*\*\**P* < 0.001, \*\*\*\**P* < 0.0001.

**Figure S5.**
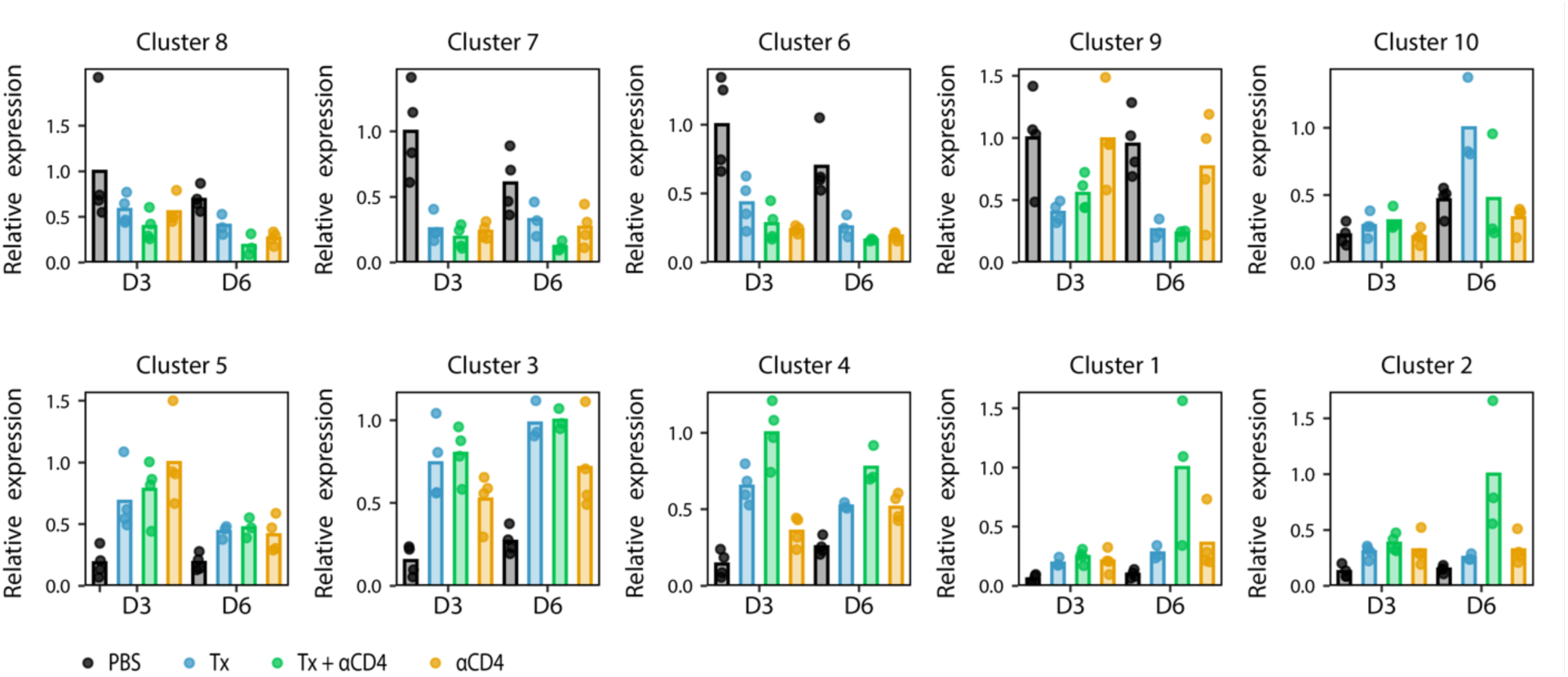
Gene clusters exhibit differential expression among various experimental cohorts. Normalized expression of all gene clusters identified in Fig 4A for different experimental cohorts.

**Figure S6.**
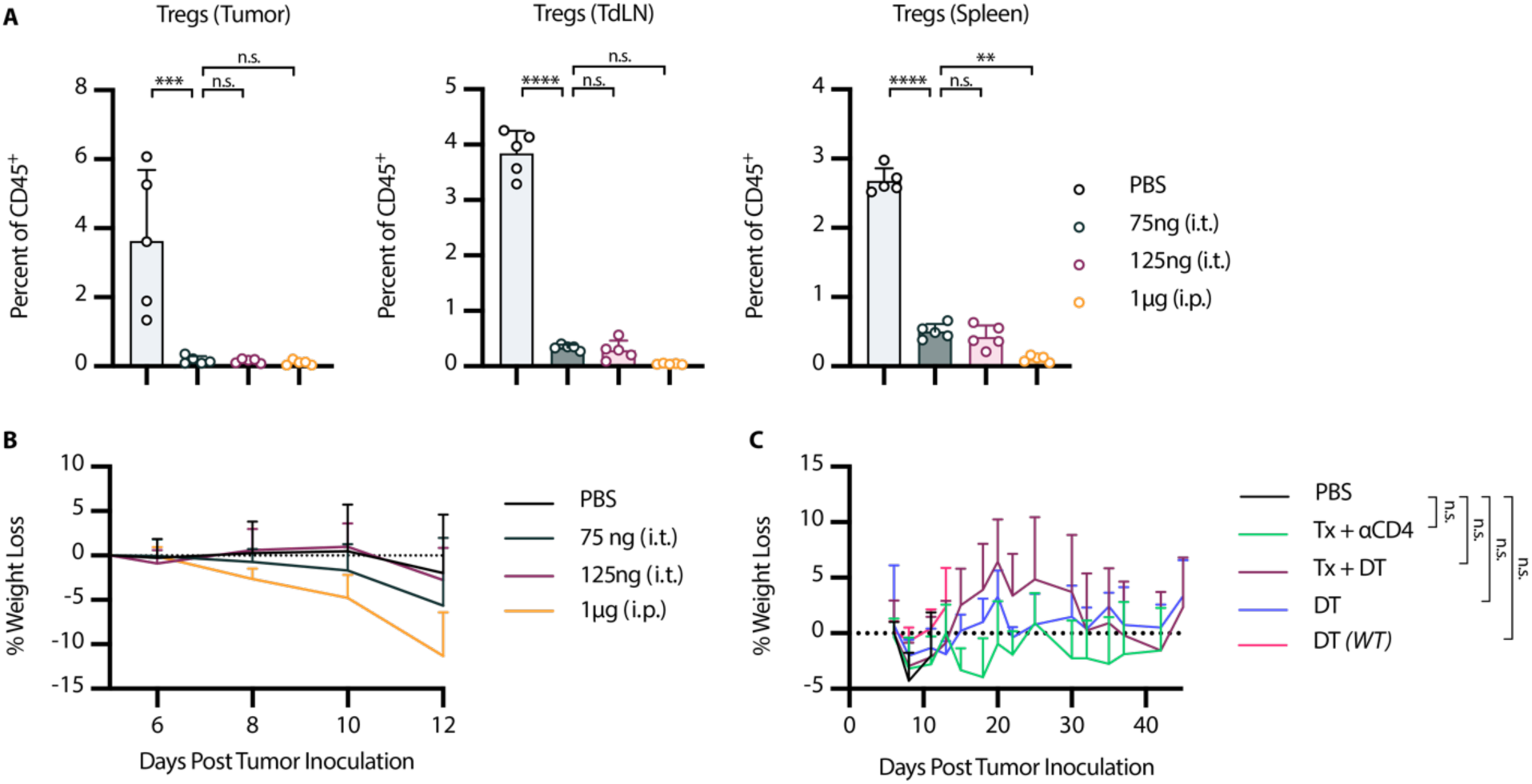
Low dose 75 ng (i.t.) DT depletes Tregs to a similar extent as 125 ng (i.t.) Foxp3-DTR Mice were inoculated with 1 x 10^6^ B16F10 cells on day 0. **(A)** Mice were treated on days 6, 8, and 10 with either 125 ng DT (i.t.), 75 ng DT (i.t.), or 1 µg DT (i.p.). Flow cytometry quantification (mean±SD) of Tregs in tumor, TdLN, or spleen on day 12 (gated on single cell/live/CD45^+^/CD3^+^NK1.1^−^/CD4^+^/GFP(*Foxp3*)^+^, n = 5) **(B)** Weight loss (mean+SD) of mice from **(A)**. **(C)** Weight loss data from survival study shown in Fig 6C. Flow data were compared using one way ANOVA with Tukey’s multiple hypothesis testing correction and weight loss data were compared using two-way ANOVA with Tukey’s multiple hypothesis testing correction. \**P* < 0.05, \*\**P* < 0.01, \*\*\**P* < 0.001, \*\*\*\**P* < 0.0001.

**Figure S7.**
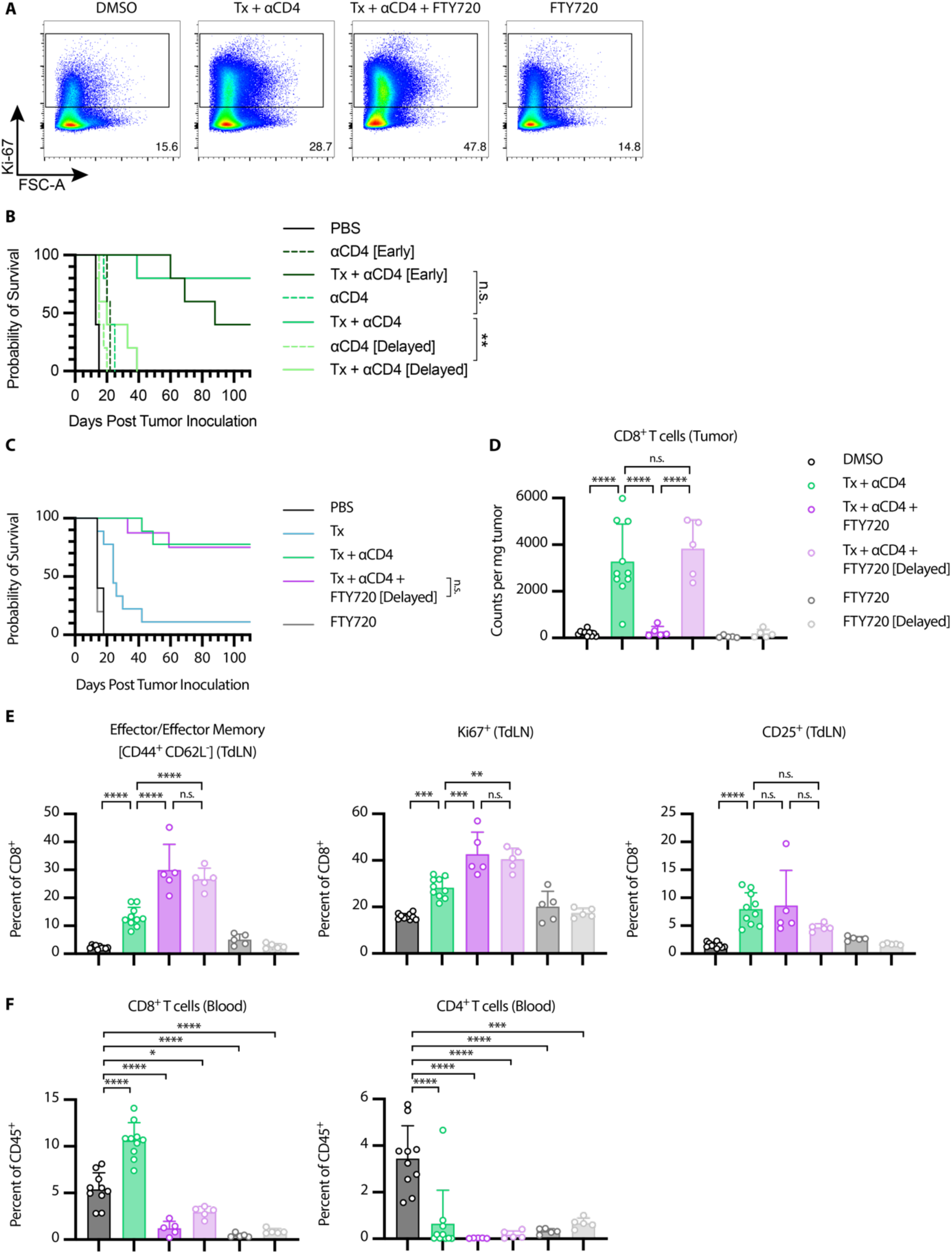
Delayed FTY720 initiation does not affect therapeutic efficacy of Tx + αCD4. **(A)** Representative gating for Ki67^+^ CD8^+^ T cells in TdLN 6 days after first α4-1BB-LAIR treatment (gated on single cell/live/CD45^+^/CD3^+^NK1.1^−^/CD8^+^). **(B-E)** Delayed FTY720 refers to FTY720 initiation concurrent with α4-1BB-LAIR treatment, while FTY720 refers to FTY720 initiation concurrent with αCD4. **(B)** Overall survival of mice treated with PBS/DMSO (n = 5), Tx (n = 9), Tx + αCD4 (n = 9), Tx + αCD4 + delayed FTY720 (n = 8), or delayed FTY720 (n = 5). Mice were treated with the same dose/dose schedule as in Fig. 1A, with delayed FTY720 treatment initiated on day 6 and continued every other day until day 34. **(C)** Flow cytometry quantification (mean±SD) of CD8^+^ T cells in the tumor 6 days after first α4-1BB-LAIR treatment (gated on single cell/live/CD45^+^/CD3^+^NK1.1^−^/CD8^+^, n = 5-10, two independent experiments). **(D)** Flow cytometry quantification (mean±SD) of effector/effector memory (CD44^+^ CD62L^−^), CD25^+^, and Ki67^+^ CD8^+^ T cells in the TdLN 6 days after first α4-1BB-LAIR treatment (gated on single cell/live/CD45^+^/CD3^+^NK1.1^−^/CD8^+^, n = 5-10, two independent experiments). **(E)** Flow cytometry quantification (mean±SD) of CD8^+^ T cells and CD4^+^ T cells in the blood 6 days after first α4-1BB-LAIR treatment (gated on single cell/live/CD45^+^/CD3^+^NK1.1^−^/CD8^+^, n = 5-10, two independent experiments). **(F)** Overall Survival of mice treated with PBS, Tx + αCD4, or αCD4 with αCD4 initiated on day 4 as outlined in figure 1A, day 10 (“delayed”), or day −8 (“early”) (n = 5). Survival data were compared using log-rank Mantel-Cox test and flow cytometry data were compared using one-way ANOVA with Tukey’s multiple hypothesis testing correction. \**P* < 0.05, \*\**P* < 0.01, \*\*\**P* < 0.001, \*\*\*\**P* < 0.0001.

**Figure S8.**
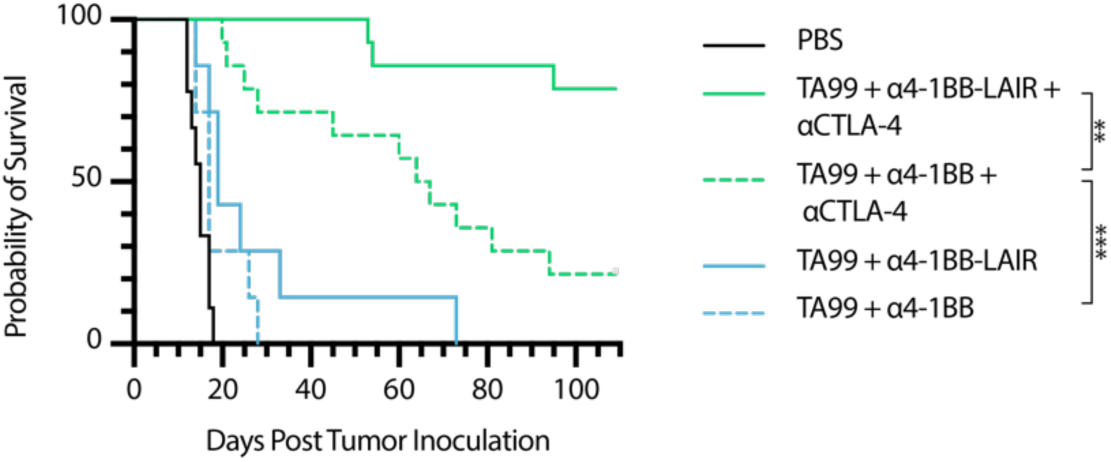
Non-collagen anchoring TA99 + α4-1BB + αCTLA-4 is also efficacious. Mice were inoculated with 1 x 10^6^ B16F10 cells on day 0. Overall survival of mice treated either with PBS (n = 9, two independent studies), Tx (n = 7), TA99 + α4-1BB-LAIR + αCTLA-4 (n = 14, two independent studies), TA99 + α4-1BB-LAIR + αCTLA-4 (n = 14, two independent studies), TA99 + α4-1BB-LAIR (n = 7), or TA99 + α4-1BB (n = 7). Mice were treated with the same dose/dose schedule as in Fig. 1A with 200 µg αCTLA-4 (i.p.) given on days 6, 9, 13, 16, 20, 23, and 27. Survival data were compared using log-rank Mantel-Cox. \**P* < 0.05, \*\**P* < 0.01, \*\*\**P* < 0.001, \*\*\*\**P* < 0.0001.

**Figure S9.**
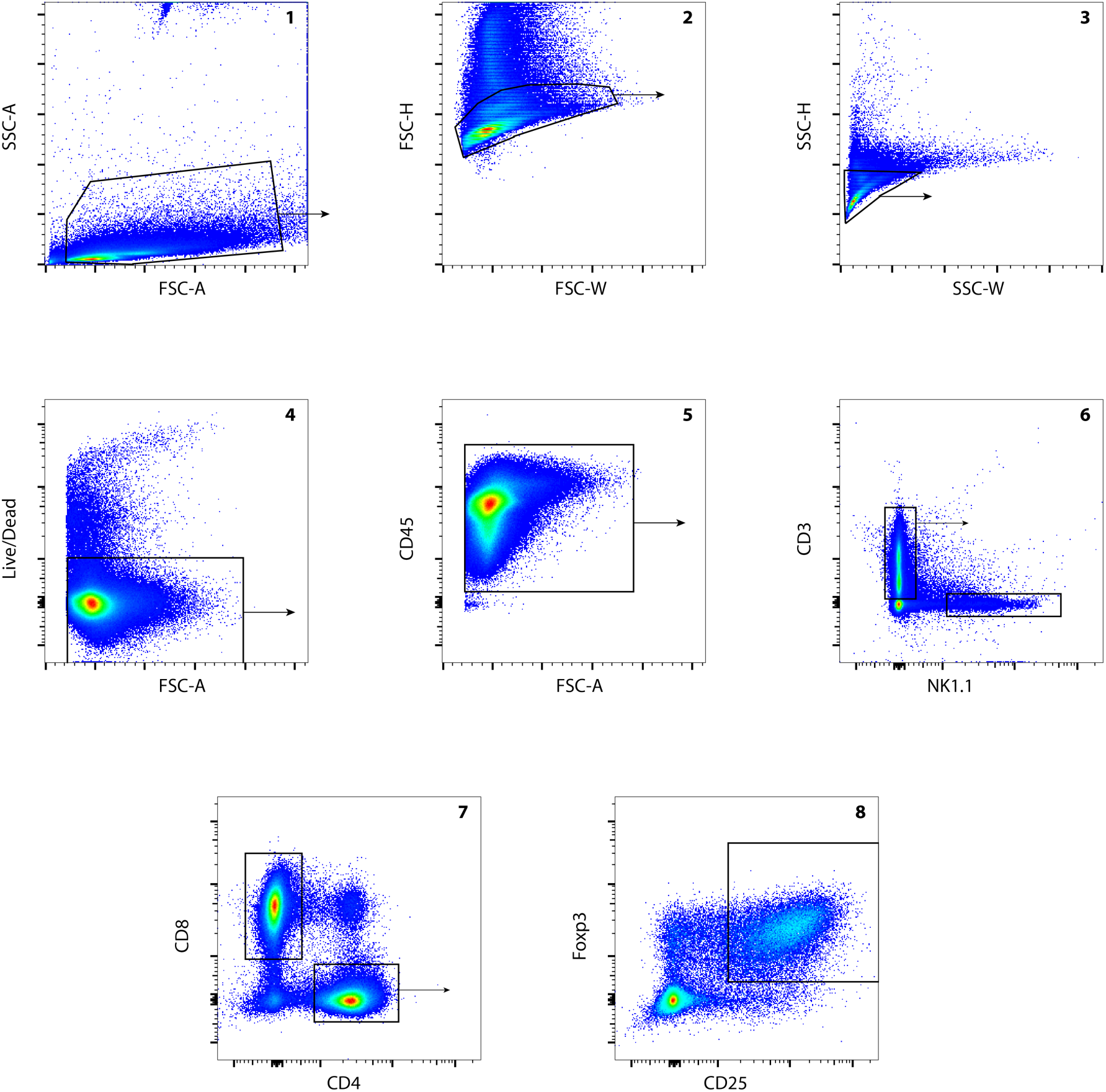
Example gating. Gating strategy for CD8^+^ T cells, CD4^+^ T cells, and Foxp3^+^ CD25^+^ Tregs, shown on a TdLN sample. Identical gating strategies were used for tumor, spleen, and blood samples.

**Figure S10.**
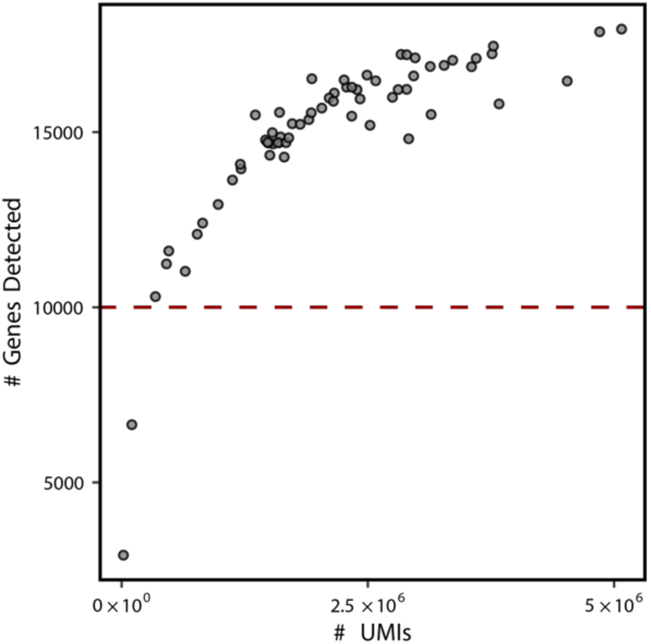
Low read samples removed from RNA-sequencing analysis. Plot of number of genes detected versus number of unique reads per sample for all tumor and TdLN bulk-RNA seq samples. Samples with less than 10,000 unique genes detected were excluded from analysis. Two samples (one Tx D6 and one Tx + αCD4 D6) met this exclusion criteria.

## SI Tables

**Table S1.**
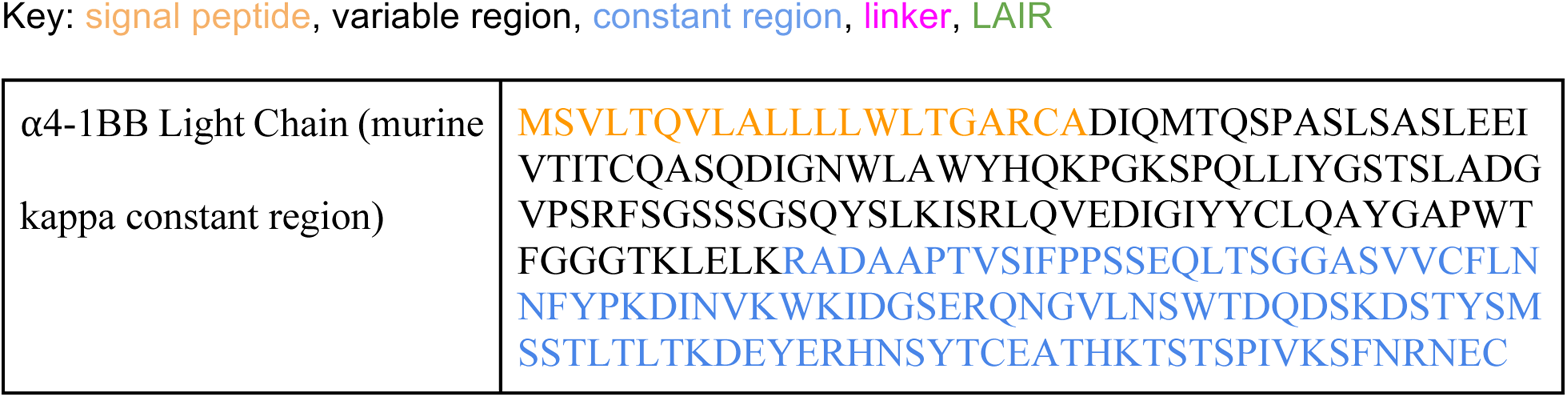
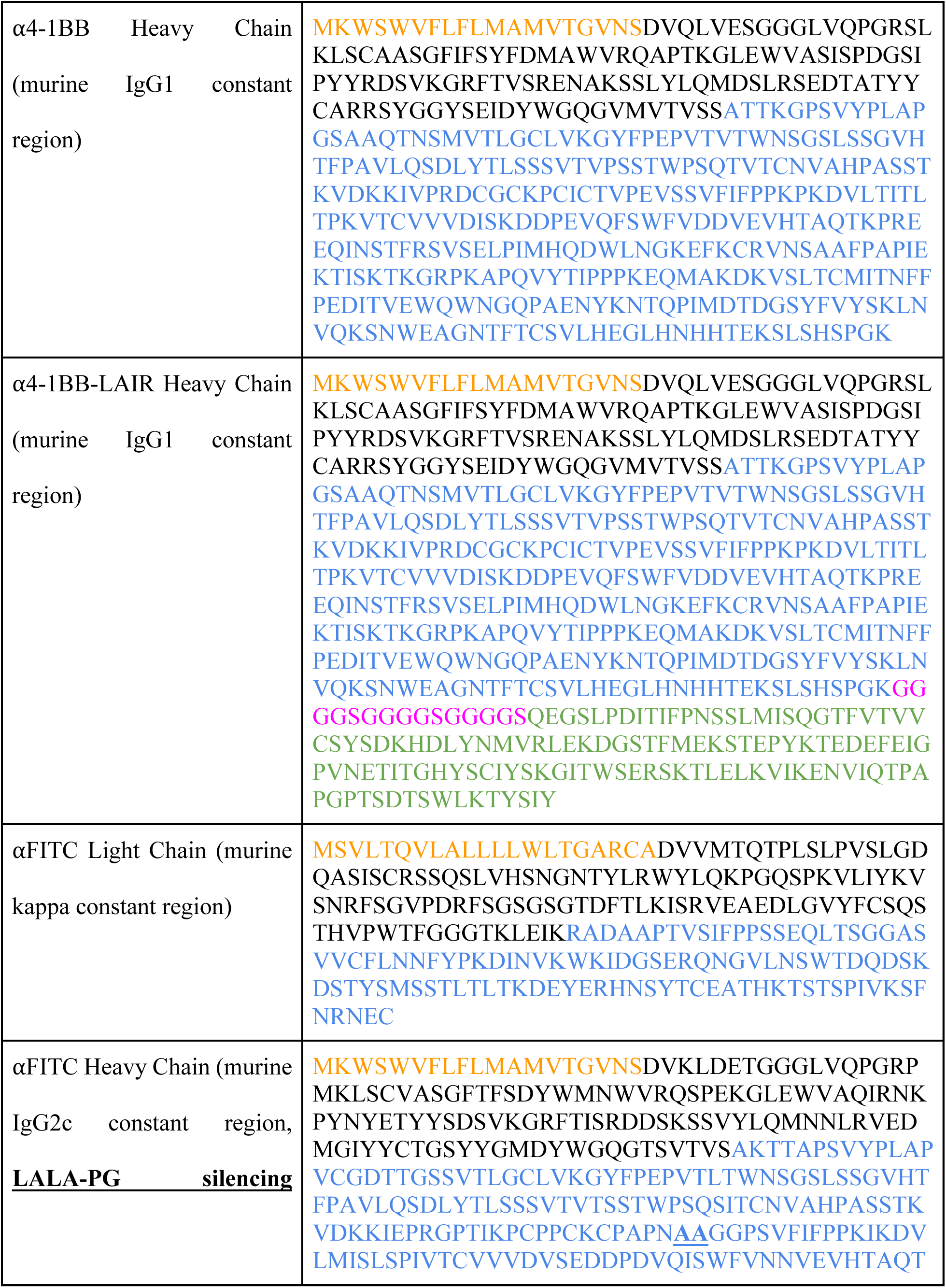
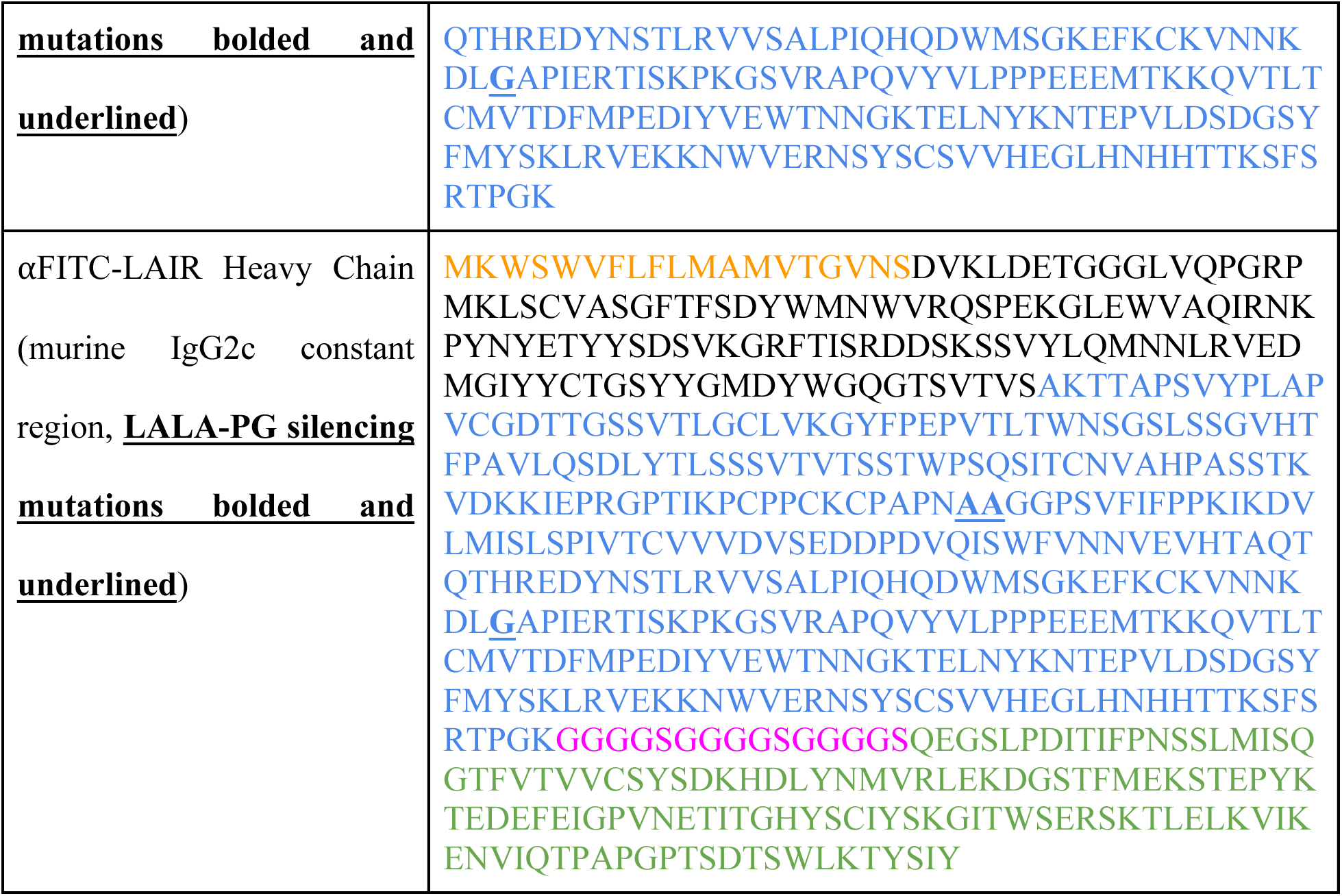
Amino acid sequence list

